# Coupling of Ca^2+^ and voltage activation in BK channels through the αB helix/voltage sensor interface

**DOI:** 10.1101/2020.01.29.925545

**Authors:** Yanyan Geng, Zengqin Deng, Guohui Zhang, Gonzalo Budelli, Alice Butler, Peng Yuan, Jianmin Cui, Lawrence Salkoff, Karl L. Magleby

## Abstract

Large conductance Ca^2+^ and voltage activated K^+^ (BK) channels control membrane excitability in many cell types. BK channels are tetrameric. Each subunit is comprised of a voltage sensor domain (VSD), a central pore gate domain, and a large cytoplasmic domain (CTD) that contains the Ca^2+^ sensors. While it is known that BK channels are activated by voltage and Ca^2+^, and that voltage and Ca^2+^ activations interact, less is known about the mechanisms involved. We now explore mechanism by examining the gating contribution of an interface formed between the VSDs and the αB helices located at the top of the CTDs. Proline mutations in the αB helix greatly decreased voltage activation while having negligible effects on gating currents. Analysis with the HCA model indicated a decreased coupling between voltage sensors and pore gate. Proline mutations decreased Ca^2+^ activation for both Ca^2+^ bowl and RCK1 Ca^2+^ sites, suggesting that both high affinity Ca^2+^ sites transduce their effect, at least in part, through the αB helix. Mg^2+^ activation was also decreased. The crystal structure of the CTD with proline mutation L390P showed a flattening of the first helical turn in the αB helix compared to WT, without other notable differences in the CTD, indicating structural change from the mutation was confined to the αB helix. These findings indicate that an intact αB helix/VSD interface is required for effective coupling of Ca^2+^ binding and voltage depolarization to pore opening, and that shared Ca^2+^ and voltage transduction pathways involving the αB helix may be involved.

**Significance:** Large conductance BK (Slo1) K^+^ channels are activated by voltage, Ca^2+^, and Mg^2+^ to modulate membrane excitability in neurons, muscle, and other cells. BK channels are of modular design, with pore-gate and voltage sensors as transmembrane domains and a large cytoplasmic domain CTD containing the Ca^2+^ sensors. Previous observations suggest that voltage and Ca^2+^ sensors interact, but less is known about this interaction and its involvement in the gating process. We show that a previously identified structural interface between the CTD and voltage sensors is required for effective activation by both voltage and Ca^2+^, suggesting that these processes may share common allosteric activation pathways. Such knowledge should help explain disease processes associated with BK channel dysfunction.

## Introduction

Large conductance Ca^2+^- and voltage-activated K^+^ channels (BK, Slo1, Maxi-K, K_Ca_1.1, *KCNMA1* gene) are widely distributed and opened by the synergistic action of depolarization and intracellular Ca^2+^ (1-17), with intracellular Mg^2+^ modulating the activation (18, 19). Open BK channels allow the outward movement of intracellular K^+^, which drives the membrane potential in a negative direction, decreasing the excitability of cells by deactivating voltage dependent Na^+^ and Ca^2+^ channels. Through this negative feedback system, BK channels modulate many physiological processes, including smooth muscle contraction (20), release of neural transmitters (21), repetitive firing in neurons (22), and renal K^+^ secretion (23). Defects in BK channels can lead to epilepsy and dyskinesia (24, 25), hypertension (23), and contribute to human obesity (26). The voltage- and Ca^2+^-dependent gating of BK channels can be described by two-tiered allosteric gating mechanisms formulated as discrete state models (5, 11), and also by more general models (6), including the modulation by Mg^2+^ (18, 27, 28). However, less is understood about the specific allosteric pathways involved in voltage and Ca^2+^ activation of the channel.

It is known that the BK channel is of modular design, with the voltage sensors and pore gate forming the transmembrane domain of the channel (TMD), and the large cytosolic domain (CTD) containing the Ca^2+^ sensors (3, 4, 7-17). Removing the CTD removes all Ca^2+^ sensitivity (29), leaving a voltage gated channel (29, 30). However, the coupling between voltage sensors and the pore gate is greatly reduced in the absence of the CTD (30), leading to a marked decrease in voltage activation (29, 30). This suggests that the CTD may also contribute to transduction from voltage sensors to the pore gate. Previous studies have identified a dynamic interface between the αB helix of the CTD and the voltage sensors (16, 17, 31). The purpose of our paper is to determine the contributions of the αB helix at this interface to voltage and Ca^2+^ activation.

The structure of one of the four identical subunits of a BK channel is shown by the cartoon in Fig. 1*A*, and the complete cryo-EM structures of metal free and Ca^2+^ and Mg^2+^ bound BK channels are shown in Figs. 1*C, D*. Each subunit has a voltage sensor domain (VSD) comprised of transmembrane segments S1-S4 that are preceded by an S0 transmembrane segment that modulates the voltage sensor (32). Following S4 are transmembrane segments S5-S6, that contribute to the central pore gate domain (PGD), which is formed from four such S5-S6 pairs, one from each subunit. The location of the gate that blocks the central pore is debated, but movement of the pore lining segment S6 in the intracellular cavity near the entrance to the selectivity filter may reposition hydrophobic residues to exclude water, blocking conduction (33, 34). The four VSDs surround the central pore gate domain.

**Fig. 1.**
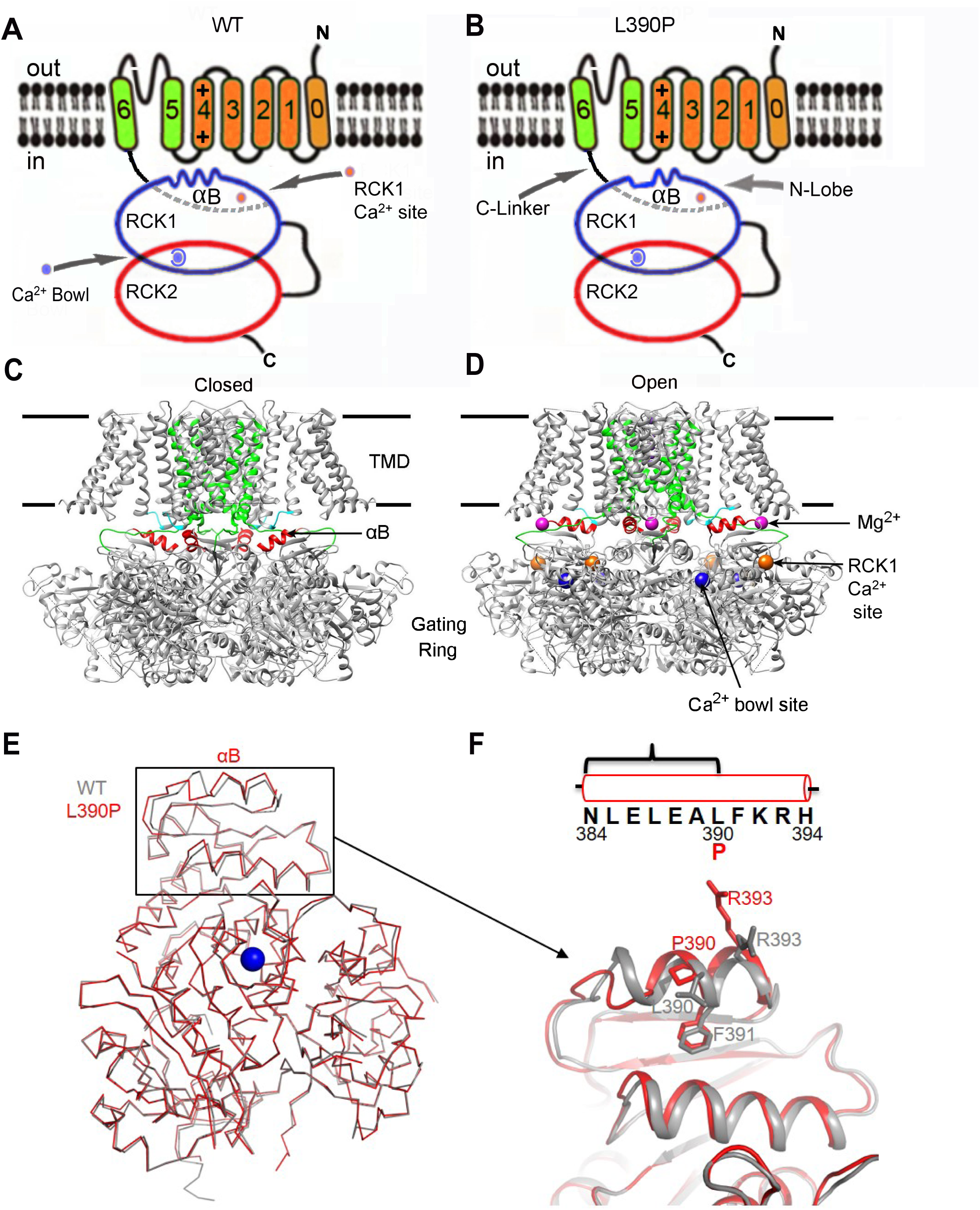
Cartoon, cryo-EM structures, and crystal structures of specified parts of the BK channel. (***A, B***) Cartoon of one subunit of the tetrameric BK channel modified from (53). In the channel the αB helix is under the S4-S5 linker/VSD of an adjacent subunit is a swapped manner. S1-S4 form the voltage sensor, S5-S6 contribute to the pore-gate domain, a C-linker connects S6 to the N-lobe of the RCK1 domain. Ca^2+^ bowl and RCK1 Ca^2+^ sites are shown. The αB helix of the N-lobe interacts with the S4-S5 linker and voltage sensor from an adjacent subunit in a swapped configuration (16). L390P disrupts the first turn of the αB helix, which would reduce the effectiveness of the αB helix/S4-S5 linker voltage sensor interface. (***C***) Cryo-EM structure of the *Aplysia* BK channel in the EDTA Ca^2+^ and Mg^2+^ free, presumably closed, state (PDB:5TJI) (17), and (***D***), in the Ca^2+^ and Mg^2+^ bound, presumably open, state PDB:5TJ6 (16). S0-S4 which include the voltage sensor are in silver, S5-S6 are in green, and the S4-S5 linkers in cyan. The αB helices (red) forming the top of the RCK1 N-lobes move upwards and outwards from closed to open, where they could push upwards and outwards on the S4-S5 linkers and voltage senosr domains. Ca^2+^ and Mg^2+^ sites are indicated as colored balls. (***E, F***) The proline substitution L390P of the αB helix caused local perturbations limited to flattening the first turn of the αB helix. (***E***) Superposition of crystal structures of WT human BK CTD in grey (PDB:3MT5) (54) and the L390P mutant CTD structure determined in this paper in red (PDB:6V5A). The most N-terminal residue is K343. Refinement details in *SI Appendix*, Table S1. (***F***) A close-up view shows that the L390P mutation flattens the first turn of the αB helix (WT in grey, mutant in red). A schematic of the residues in the αB helix is above the structure of the helix. The bracket includes the first residue with altered position N384 and ends with L390P.

The large cytosolic domain (CTD) of the BK channel, also referred to as the gating ring (31, 35) is seen in Fig. 1*C, D*). Each subunit contributes tandem RCK1 and RCK2 domains to the CTD, for a total of eight RCK domains. Each RCK1 contains a high affinity RCK1 Ca^2+^ binding site (7, 36), and each RCK2 contains a high affinity Ca^2+^ bowl binding site (4), for a total of eight high affinity Ca^2+^ sites in the CTD. In addition, there are four low affinity Mg^2+^ sites per channel, each coordinated between residues on the top of the CTD and the bottom of the voltage sensor (28). Removing the CTD removes all Ca^2+^ and Mg^2+^ sensitivity (29).

The CTD is anchored to the TMD through four peptide C-linkers, each connecting S6 of the TMD to the N-lobe of the RCK1 domain of the same subunit in a swapped manner. BK channels have a relatively short S4-S5 linker extending from S4 in the VSD to S5 in the PGD of the same subunit. The short S4-S5 linker in BK would constrain the depolarization induced upward movement of S4 in BK (16, 17), which may explain why only one positive residue in each S4 contributes to the voltage dependence of gating in BK (37).

Ca^2+^ binding to the Ca^2+^ bowl and RCK1 sites in the CTD leads to a near rigid-body lateral tilting (away from pore) of the N-lobes formed by the upper part of each RCK1 domain (16, 17, 31). This lateral tilting moves the RCK1 attachment point for each S6-RCK1 C-linker laterally and downwards, pulling on S6 to potentially open the pore gate. Simultaneously, the lateral tilting of the N-lobe moves the αB helix located at the top of each N-lobe both upwards and laterally to push upwards and laterally on the bottom of S4-S5 linker/VSD to potentially open the channel (16, 17). A morphing movie in Hite, Tao, and MacKinnon (17) clearly illustrates these simultaneous actions of the Ca^2+^-induced pulling on S6 through the C-linker and the upwards and lateral pushing on the S4-S5 linker/VSD. A similar upward and outward movement of the αB helices at the top of the N-lobes induced by 10 mM Ca^2+^ for vertebrate BK channels can be seen in a movie in Yuan et al. (31). Supporting that Ca^2+^-induced pulling on the S6-RCK1 linkers contributes to channel opening are the observations that shortening the C-linkers by deleting amino acids increases open probability over a range of Ca^2+^ concentrations, and that lengthening the C-linkers decreases open probability (38, 39).

In the absence of Ca^2+^, shortening the C-linkers left shifts V_1/2_, the voltage for half activation, to more negative potentials, and lengthening the C-linkers right shifts V_1/2_ to more positive potentials (39). These shifts in voltage activation induced by changing the length of the C-linker arising from the Ca^2+^-sensing CTD, suggests coupling between the Ca^2+^ and voltage activation pathways. This observation is one of many suggesting coupling between voltage and Ca^2+^activation in BK channels, as detailed in reviews (14, 15, 40). A few of these will be mentioned. Removing the CTD right shifts V_1/2_ by > 50 mV (29) by decreasing the coupling between voltage sensor movement and the pore gate (30). A small but significant interaction between Ca^2+^ binding and voltage sensor activation is required to account for the Ca^2+^ dependence of Po (6). Ca^2+^ binding to the RCK1 Ca^2+^ site is voltage dependent (41). Functional studies suggest that residues in the AC region (N-lobe) of the CTD interact with the VSD (25). Ca^2+^ binding alters VSD rearrangement (42). Ca^2+^ sensor occupancy left shifts the Q-V curves, with strong allosteric coupling between Ca^2+^-binding sites and voltage sensors (43). Strong correlations have been observed between VSD rearrangements and conformational changes in the CTD, which support a pathway that couples Ca^2+^ binding to channel opening through the voltage sensor (44). Mg^2+^ binding between the VSD and the CTD directly couples voltage, Ca^2+^, and Mg^2+^ sensors (19, 28).

The structural studies (16, 17) discussed above and Fig. 1*C, D*, showing that the αB helices at the top of the N-lobe of the CTD interact directly with the S4-S5 linker/VSD, implicate the αB helix as a potential key structural element in the coupling between voltage and Ca^2+^ sensors and suggest that the αB helix may be part of a shared pathway for voltage and Ca^2+^ activation (17). Functional support for this proposal is the finding from an alanine scan of the AC region of the N-lobe, that alanine substition at two sites in the αB helix decreased both Ca^2+^ sensitivity (25) and interactions among residues in the αB helix and VSD (19). A prediction of coupling of voltage and Ca^2+^ sensors through the αB helix is that disruption of the αB helix should alter both Ca^2+^ and voltage activation of the BK channel.

In this paper we test this hypothesis by disrupting the αB helix with point proline mutations. We first examined a proline substitution in the middle of the helix, and found that L390P shifted voltage activation to more positive potentials and decreased Ca^2+^ sensitivity. A high resolution crystal structure of the CTD L390P mutant showed that the effects of the mutation were local to the αB helix, limited to flattening one turn of the αB helix. L390P did not alter the normalized gating currents. Analysis of Po versus voltage data with the HCA model (45) indicated that L390P reduced voltage activation by decreasing the coupling between voltage sensors and the pore gate. Mutations eliminating either the high affinity RCK1 Ca^2+^ sites or the Ca^2+^-bowl sites showed that L390P still reduced Ca^2+^ activation when either type of site was examined separately, suggesting that both types of high affinity Ca^2+^ sites require an intact αB helix for full function. Proline mutations at other sites in the αB helix also reduced both voltage and Ca^2+^ activation. αB helix mutations also decreased activation by 10 mM Mg^2+^ acting at the low affinity Mg^2+^ site. Our findings suggest that an intact αB helix is required for effective voltage, Ca^2+^, and Mg^2+^ activation of BK channels. Coupling of voltage, Ca^2+^, and Mg^2+^ activation through the αB helix would allow voltage, Ca^2+^, and Mg^2+^ to share activation pathways.

## Results

### Disruption of the αB Helix Right Shifts V_1/2_ to More Positive Potentials, with Little Effect on the G-V Slopes

If the αB helix plays a major role in the transduction of voltage- and Ca^2+^-activation to the pore gate in BK channels, then disruption of the αB helix might be expected to alter voltage and Ca^2+^ activation of the channel. To test this hypothesis a proline was substituted for a leucine in the middle of the three turn αB helix that interfaces with the S4-S5 linker/VSD to obtain L390P channels (Fig. 1*A, B*). X-ray crystallography then indicated that the L390P mutation caused a structural change to the αB helix by flattening the first turn of the helix, with structural changes limited to the αB helix (Fig. 1*B, E, F*).

To test if this structural changes to the αB helix altered voltage and Ca^2+^ activation, the voltage required for half-activation of the channel, V_1/2_, was determined for L390P channels for comparison to WT channels. V_1/2_ for WT and L390P channels was first obtained in the absence of Ca^2+^ to determine the effect of αB helix disruption on voltage activation. V_1/2_ was then determined in the presence of 100 µM Ca^2+^ to measure the effect of αB helix disruption on Ca^2+^ activation.

For WT channels in 0 µM Ca^2+^, the mean voltage for half activation was 176 mV (SEMs and *p* values are in *SI Appendix*, Table S2). Disruption of the αB helix with L390P then right shifted the G-V curve to more positive potentials, giving a mean V_1/2_ of 242 mV. This indicated that a 66 mV greater depolarization was required to activate the channel in the absence of Ca^2+^ after disrupting the αB helix (Fig. 2 and *SI Appendix*, Fig. S1 and Table S2). Proline substitutions at other positions along the αB helix (L385P, L387P, E388P, A389P, F391P, and F394P) also right shifted the voltage required for half activation in the absence of Ca^2+^ by from 44 to 109 mV, depending on the location of the mutation (Fig. 2 and *SI Appendix*, Fig. S1 and Table S2). Hence, more positive potentials are required to activate the channel when the αB helix is disrupted, indicating that an intact αB helix is required for effective voltage activation.

**Fig. 2.**
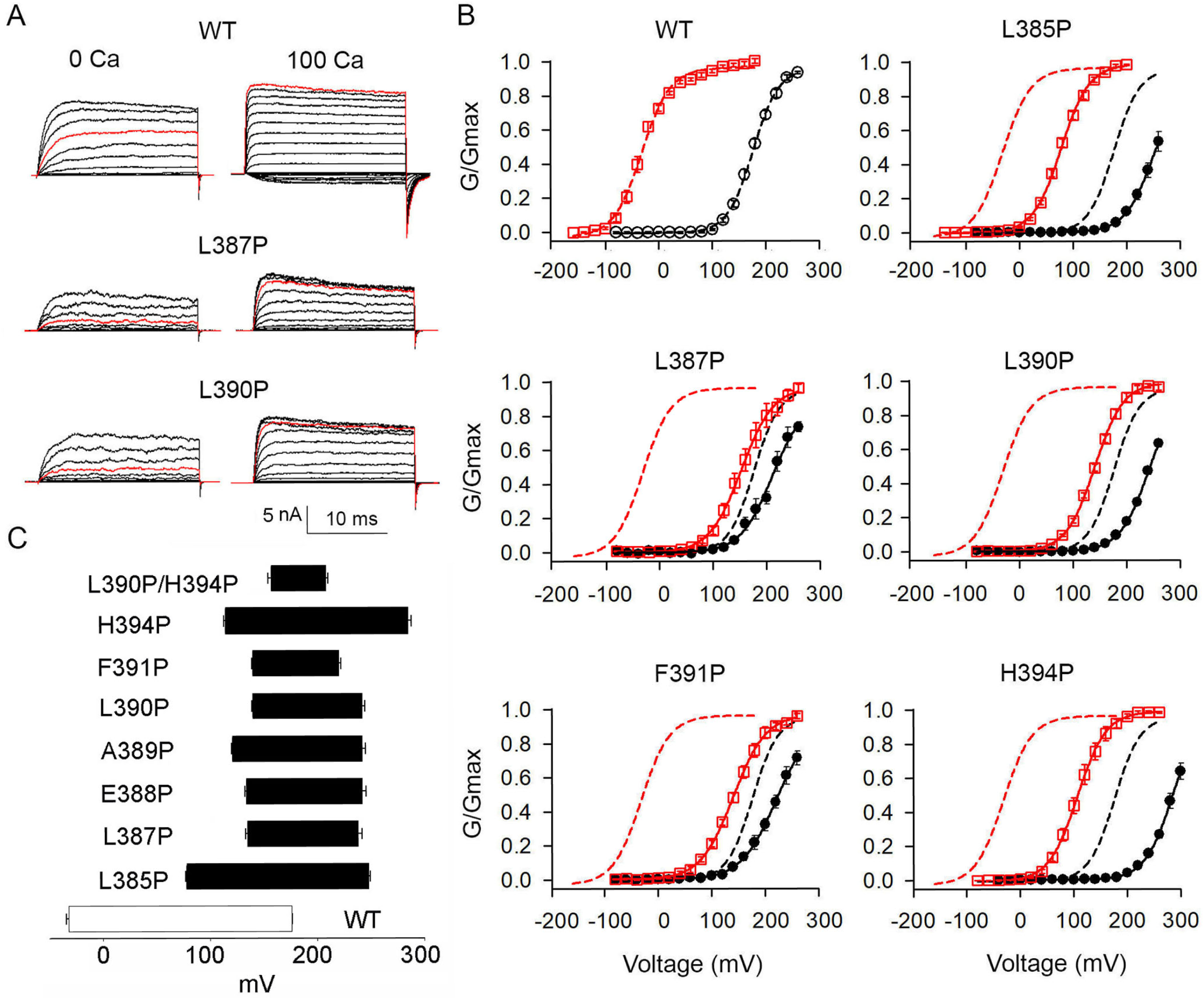
Proline mutations of the αB helix decrease both voltage and Ca^2+^ activation. (***A***) Current recordings for the indicated channels from inside-out macro patches. Voltage pulses were from - 80 to 260 mV with 20 mV increments, except for WT 100 µM Ca^2+^ which were from -160 mV to 180 mV. Red trace for WT 100µM Ca^2+^ is current at 180 mV. Other red traces are currents at 200 mV. (***B***) G/G_max_ vs. V plots in 0 Ca^2+^ (black lines through black filled circles, n≥4) and 100 µM Ca^2+^ (red lines through red open squares, n≥4). Dashed lines are Boltzmann fits for WT channels and continuous lines for mutated channels. Additional data are in Fig. S1. Boltzman fit parameters, SEMs, and *p* values are in *SI Appendix*, Table S2. (***C***) V_1/2_-Ca^2+^ bar plots, where the right end of each bar indicates 0 µM Ca^2+^ and the left end 100 µM Ca^2+^ as mean and SEM. Bar length is taken as a measure of Ca^2+^ sensitivity.

### Quantifying the protein interaction interface between the αB helix and the S4-S5 linker/VSD

Fig. 1 shows that the structural changes from mutating the αB helix with L390P were localized to the αB helix (Fig. 1*E, F*). Consequently, for the mutated αB helix to alter gating, the mutation would have to alter a functional interaction at an interface between the αB helix and some other part of the channel. Hite et al. (17) have identified a protein-protein interface between the RCK1 N-lobe and the S4-S5 linker/VSD in the open BK channel. To further quantify interface interactions, we measured the protein interaction interface (buried surface area), which is a direct measurement of protein-protein interaction, between the αB helix and the S4-S5 linker/VSD for both closed and open Aplysia BK channels. For closed channels the area per subunit of the protein interaction interface between the αB helix and the S4-S5 linker was 280 Å^2^, and between the αB helix and the combined S4-S5 linker/VSD was 299 Å^2^. For ligand bound open channels these numbers were 270 Å^2^ and 643 Å^2^, respectively, showing protein-protein interaction at the interface in both the closed and open states, with a large increase in the total protein interaction interface when the channel is open.

Our observations that the L390P mutation flattens the first turn of the αB helix, with no significant changes in structure elsewhere in the RCK1-RCK2 domains (Fig. 1*E, F*), requires that the L390P mutation alters interaction at the interface between the αB helix and the S4-S5 linker/VSD. That both Ca^2+^ and voltage activation are reduced by L390P acting at this interface suggests that this interface could provide a common pathway through which voltage and Ca^2+^ activation could interact.

### Disruption of the αB Helix with L390P has Negligible Effect on Gating Charge Movement

In contrast to the large right shifts in V_1/2_ required for activation with proline substitution in the αB helix, the voltage sensitivity of activation, as indicated by the slopes of the G-V curves, remained relatively unchanged (Fig. 2 and *SI Appendix*, Fig. S1 and Table S2). Similar slopes suggest similar effective charge movement for activation.

If effective charge movement is similar after disruption of the αB helix, what then accounts for the large right shifts in V_1/2_? One possibility is that the movement of the voltage sensors occurs at right shifted voltages after disrupting the αB helix. Another is that pore opening is no longer tightly coupled to voltage sensor movement (30). A comparison of gating currents recorded from WT and L390P channels indicated that disruption of the αB helices had negligible effect on Q-V curves (Fig. 3). Hence, the right shift in the G-V curves with L390P does not arise from a right shift in voltage sensor activation.

**Fig. 3.**
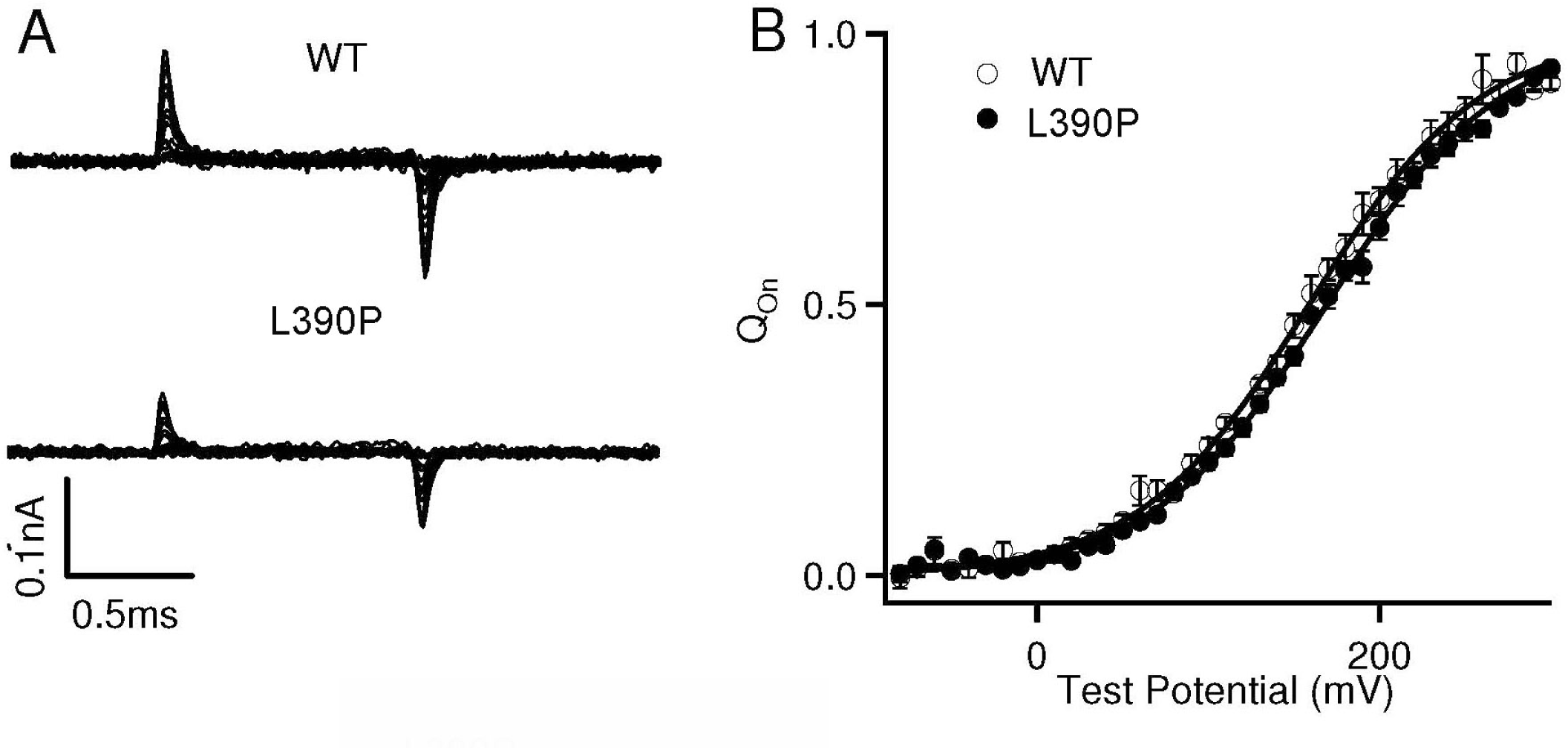
Disrupting the αB helix with L390P has negligible effect on the gating current. (***A***) Gating currents of WT and L390P BK channels. Voltage pulses were from -30 to 300 mV (WT) or from - 80 to 300 mV (L390P) with 20 mV increments. (***B***) Normalized gating charge-voltage (Q-V) curves for on-gating currents. The smooth curves are fits to the Boltzmann function with a V_1/2_ and slope factor of 159.1 ± 6.5 mV and 49.0 ± 6.7 mV for WT, and 169.0 ± 6.7 mV and 50.7 ± 5.1 mV for L390P. The data points represent the mean ± SEM. n=8 for WT and n=10 for L390P.

### Disruption of the αB Helix with L390P Increases the Intrinsic C-O Equilibrium Constant *L*_0_ and Decreases the Coupling Factor *D* between Voltage Sensors and Pore Gate

We next considered whether the L390P induced right shift in G-V curves arose from a decrease in the intrinsic closed-open equilibrium constant *L*_0_ or from a decrease in the allosteric factor *D* that couples voltage sensor activation to pore gate opening (30, 45). Decreasing either of these factors would make it more difficult to open the channel. Po was measured over wide ranges of voltage for WT and L390P channels and the data fitted with the HCA (45) model to estimate *L*_0_ and *D.* Compared to WT, L390P increased Po at negative potentials where the channels opened spontaneously with the voltage sensors nearly at rest (Fig. 4*B, C*), and decreased Po at positive potentials (Fig. 4*C*). Fitting the Po versus V curve in Fig. 4 with the HCA (45) model indicated that L390P significantly increased *L*_0_ 12.8 fold, from 7.42×10^−7^ ± 0.96×10^−7^ for WT channels to 9.47×10^−6^ ± 2.17×10^−6^ for L390P channels (*p*<0.01). L390P also significantly decreased the coupling factor *D* by 4.94 fold, from 29.3 ± 3.0 for WT channels to 5.93 ± 0.73 for L390P channels (*p*< 0.0001).

**Fig. 4.**
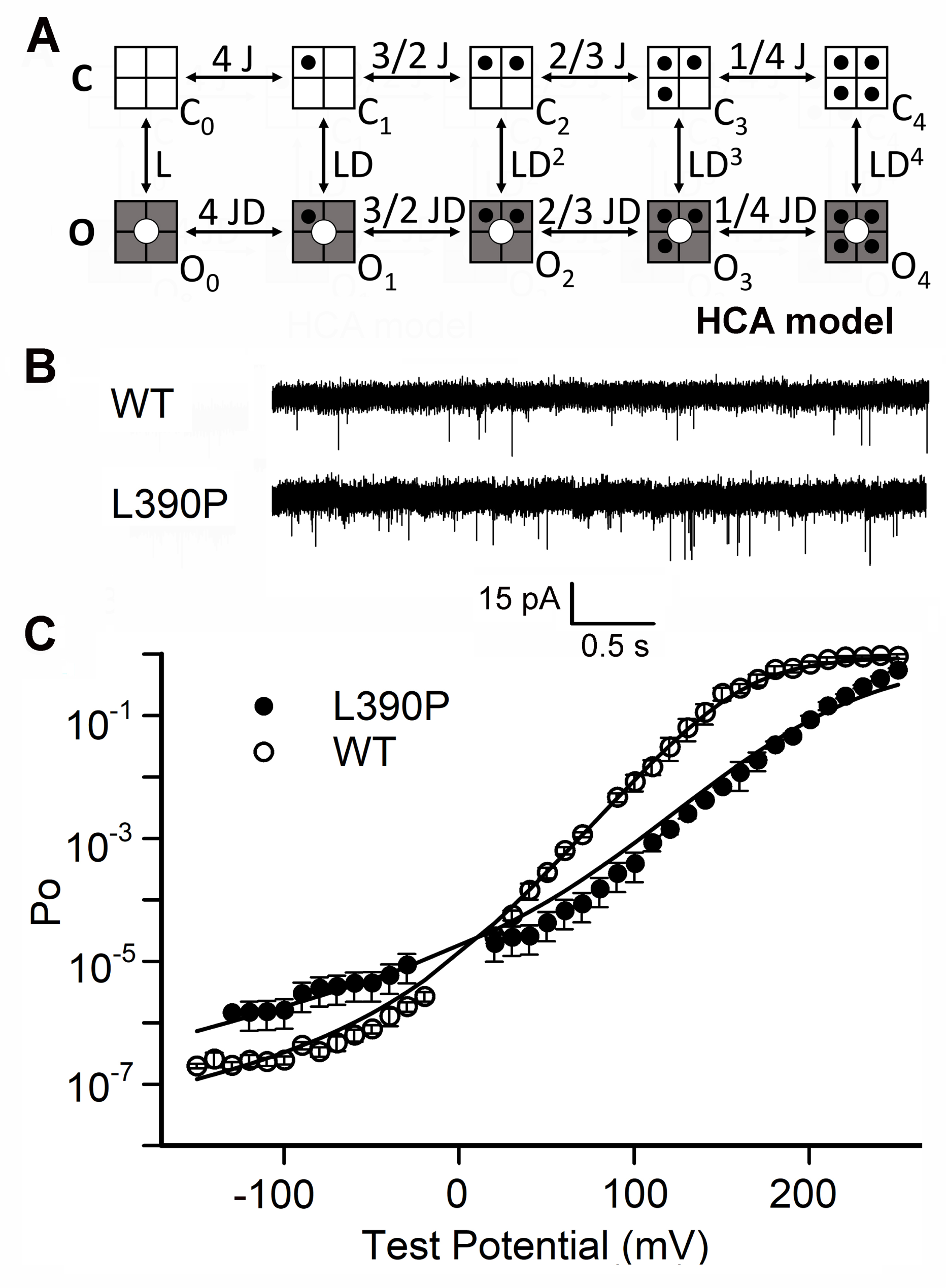
L390P increases the intrinsic closed-open equilibrium and decreases the coupling between voltage sensors and pore-gate. (***A***) HCA model (45) for voltage dependent gating of BK channels. *L* is the closed to open equilibrium constant for pore opening. *D* is the allosteric factor coupling voltage sensor activation to channel opening. *J* is the relaxed to activated equilibrium constant for each voltage sensor when the pore-gate is closed. Channel opening increases *D*-fold for each activated voltage sensor (black dot). *L* and *J* increase with depolarization. (***B***) Single channel currents recorded at -120 mV from membrane patches containing many channels. (***C***) Plot of Po versus voltage for WT and L390P channels. WT data from Zhang et al. (30). The continuous lines are fits with the HCA (45) model with parameters for WT: *L*_0_ = 7.42E-7, *D* = 29.3, *Z*_J_ = 0.520 e_o,_ *Z*_L_= 0.341 e_o_, Vhc = 159 mV, and for L390P: *L*_0_ = 9.47E-6, *D* = 5.94, *Z*_J_ = 0.503 e_o,_ *Z*_L_ = 0.440 e_o_, Vhc = 169 mV, where *L*_0_ is *L* at 0 mV. n≥4.

In terms of the HCA model, the L390P induced increase in Po at negative voltages arose from a 12.8 fold increase in the intrinsic closed-open equilibrium constant *L*_0_, and the decrease in Po at positive voltages arose from a 4.94 fold decrease in the coupling factor *D* between voltage sensors and pore gate. An increase in *L*_0_ in isolation would left shift G-V curves. Consequently, the observed increase in *L*_0_ is not responsible for the right shift at positive potentials observed in Fig. 2. The right shift occurs because the 4.94 decrease in coupling factor *D* is applied at increasing power with each activated voltage sensor (Fig. 4*A*). With four activated voltage sensors, the L390P mutation decreases effective coupling by 29.3^4^(WT)/5.93^4^(L390P) or 596 fold. The increased *L*_0_ suggests that disruption of the αB helix removes a normal inhibition on opening in WT channels. Similar to the dual action of the L390P mutation on increasing *L*_0_ and decreasing *D*, removing the entire gating ring also removes a normal inhibition on opening and decreases coupling between voltage sensors and pore gate (30).

### Disruption of the αB Helix Decreases High Affinity Ca^2+^ Sensitivity

Removing the CTD (29) or mutating both the RCK1 and calcium bowl Ca^2+^ binding sites in the CTD (7, 8) removes all Ca^2+^ sensitivity from BK channels. Hence, the CTD is necessary to both bind Ca^2+^ and to transfer the energy of Ca^2+^ binding in the CTD to the PGD to open the channel. If the αB helix is involved in this transduction, then disruption of the αB helix might be expected to reduce Ca^2+^ sensitivity. This question was explored by using the Ca^2+^-induced leftward shift in V_1/2_ as a measure of Ca^2+^ sensitivity (46). For WT BK channels, increasing Ca^2+^ from 0 to 100 µM left shifted V_1/2_ from 176 mV to -32 mV, for a Δ shift of -208 mV (Fig. 2 and *SI Appendix*, Table S2 which includes SEMs and *p* values). Thus, for WT channels, 0 to 100 µM Ca^2+^ provided activation equivalent to Δ208 mV of depolarization, as the voltage had to be shifted Δ-208 mV to negate the opening effect of 100 µM Ca^2+^.

In contrast, after disrupting the αB helix with L390P, increasing Ca^2+^ from 0 to 100 µM left shifted V_1/2_ from 242 mV to 140 mV, for a Δ shift of -102 mV, only 49% of the Δ-208 mV Ca^2+^-induced left shift for WT (Fig. 2 and *SI Appendix* Table S2). Thus, disrupting the αB helix with L390P reduced the effective transduction of high affinity Ca^2+^ binding from the gating ring to the pore gate by 51%.

Proline substitutions at positions L385P, L387P, E388P, A389P, F391P, and H394P in the αB helix also reduced the Ca^2+^-induced left shift in V_1/2_, with reductions to 82%, 50%, 52%, 58%, 38%, and 82% of WT, respectively (Fig. 2 and *SI Appendix*, Fig. S1 and Table S2). L387P and F391P gave the greatest reduction in Ca^2+^ sensitivity, consistent with previous observations obtained from an alanine scan of the αB helix, where L387A and F391A were found to reduce Ca^2+^ sensitivity (25). The reduction in Ca^2+^ sensitivity by L387A and F391A was less than half the reduction with proline substitutions at these sites, indicating that side-chain replacement with alanine can also reduce Ca^2+^ sensitivity, but that disrupting the αB helix with proline is more effective. Taken together, the observations in this section indicate that an intact αB helix is required to fully couple Ca^2+^ binding in the CTD to pore opening in the PGD.

Interestingly, the proline mutations L385P and H394P near the ends of the αB helix gave the smallest reductions in Ca^2+^ sensitivity but the greatest right shifts of V_1/2_ for voltage activation (Fig. 2 and *SI Appendix*, Fig. S1 and Table S2), indicating differential effects of the αB helix on Ca^2+^ and voltage activation, depending on the position of the substituted residues.

### Disruption of the αB Helix Reduces High Affinity Ca^2+^ Sensitivity through Both the Ca^2+^ Bowl and RCK1 Ca^2+^ Sites

The previous section showed that disrupting the αB helix with L390P and other proline mutations reduced Ca^2+^ activation. To examine whether this reduction arose from decreased effectiveness of one or both high affinity Ca^2+^ sensor pathways, we tested the effect of disrupting the αB helix on each Ca^2+^ sensor separately. The mutation D898-902N (5D5N) (4, 7) was used to eliminate the Ca^2+^ bowl, leaving only the RCK1 Ca^2+^ sensor pathway, and the mutation D362A/D367A (2D2A) (7) was used to eliminate the RCK1 Ca^2+^ site, leaving only the Ca^2+^ bowl sensor pathway.

ΔV_1/2 0→100 µM Ca_ was -208 mV for WT channels, -166 mV for 5D5N channels, and -54 mV for L390P/5D5N channels (Fig. 5 and *SI Appendix*, Fig. S2 and Table S3 which includes SEMs and *p* values). Removing the Ca^2+^ bowls thus reduced Ca^2+^ sensitivity to 80% of WT, indicating that RCK1 Ca^2+^ sites in the absence of Ca^2+^ bowls can contribute 80% of the total Ca^2+^ sensitivity of WT channels in the absence of Ca^2+^ bowls. Disrupting the αB helix of these 5D5N channels with L390P (L390P/5D5N channels) then reduced Ca^2+^ sensitivity to 33% of 5D5N channels (Fig. 5 and *SI Appendix*, Fig. S2 and Table S3).

**Fig. 5.**
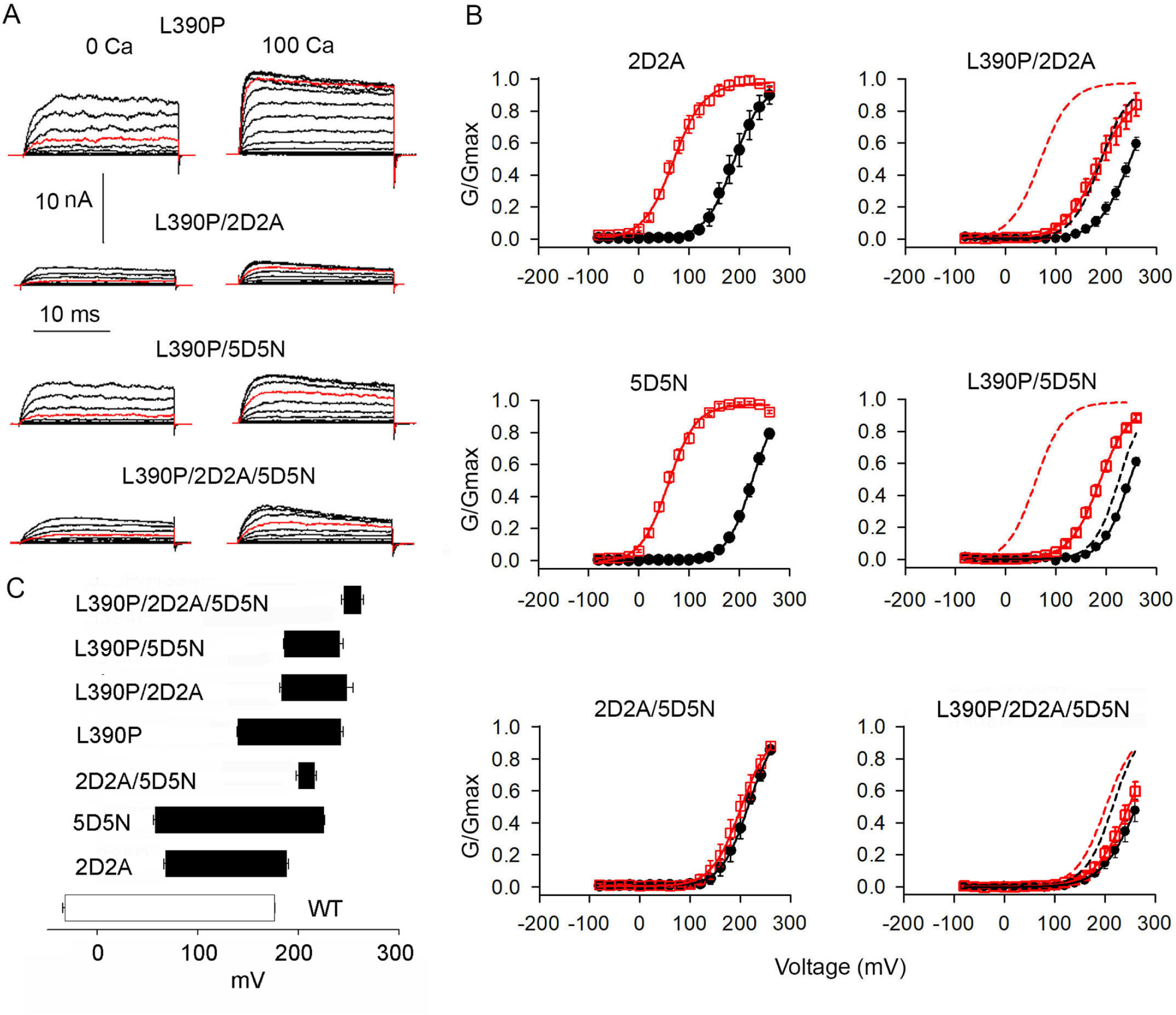
Both the Ca^2+^ bowl and RCK1 Ca^2+^ binding sites require an intact αB helix for effective Ca^2+^ activation. (***A***) Current recordings from the indicated channels. Red traces are currents at 200 mV. Voltage pulses were from -80 to 260 mV with 20 mV increments. 2D2A = D362A/D367A, disrupts the RCK1 Ca^2+^ binding site and 5D5N = D(898-902)A disrupts the Ca^2+^ bowl binding site (7). (***B***) G/G_max_ vs. V plots in 0 Ca^2+^ (black lines through black filled circles, n≥4) and 100 µM Ca^2+^ (red lines through red filled circles, n≥4). Dashed lines are Boltzmann fits for WT channels and continuous lines for mutated channels. Additional data in Fig. S2, with Boltzmann fit parameters, SEMs, and *p* values are in *SI Appendix*, Table S3. (***C***) V_1/2_-Ca^2+^ bar plots, where the right end of each bar indicates 0 µM Ca^2+^ and the left end 100 µM Ca^2+^, as mean and SEM. n≥4.

ΔV_1/2 0→100 µM Ca_ was -208 mV for WT channels, -119 mV for 2D2A channels, and -64 mV for L390P/2D2A channels. Removing the RCK1 Ca^2+^ sites thus reduced Ca^2+^ sensitivity to 57% of WT, indicating that the Ca^2+^ bowls in the absence of RCK1 sites can contribute 57% of the total Ca^2+^ sensitivity of WT channels. Disrupting the αB helix of these 2D2A channels with L390P (L390P/2D2A channels) then reduced the Ca^2+^ sensitivity to 54% of 2D2A channels. Thus, both types of high affinity Ca^2+^ sites, Ca^2+^ bowl and RCK1, require an intact αB helix for their Ca^2+^ activation to be fully functional. However, L390P had differential effects on reducing the Ca^2+^ sensitivity from the two Ca^2+^ sites. L390P/5D5N channels had 33% of the Ca^2+^ sensitivity of 5D5N channels, whereas L390P/2D2A channels had 54% of the Ca^2+^ sensitivity of 2D2A channels (Fig. 5 and *SI Appendix*, Fig. S2 and Table S3).

In the absence of L390P mutations, our observations that RCK1 Ca^2+^ sites acting alone make a greater contribution to Ca^2+^ activation than Ca^2+^ bowl sites acting alone, and that the sum of ΔV_1/2 0→100 µM Ca_ for RCK1 Ca^2+^ sites alone and Ca^2+^ bowl sites alone can exceed ΔV_1/2 0→100 µM Ca_ for WT channels has been described previously (47, 48). Also observed by others have been equal contributions of the RCK1 Ca^2+^ site and Ca^2+^ bowl to left shifts in V_1/2_ (7, 8, 49), and equal contributions of these sites to decreasing the free energy necessary to activate voltage sensors (43). The reason for the apparent differences in the relative contributions from the two types of Ca^2+^ sites is not known.

### Disruption of the αB Helix Reduces Low Affinity Mg^2+^ Activation

The previous sections showed that disruption of the αB helix reduced high affinity Ca^2+^ activation through both Ca^2+^ bowl and RCK1 Ca^2+^ sites. To explore whether proline mutations of the αB helix also changed activation through the low affinity Mg^2+^ site, we compared the shift in V_1/2_ for mutated and WT channels for 0 →10 mM Mg^2+^. In these experiments, Mg^2+^ would be expected to act predominantly on the low affinity Mg^2+^ sites (27). Results for Mg^2+^ activation of WT channels and for seven different proline substitutions in the αB helix are presented (Fig. 6 and *SI Appendix*, Fig. S3 and Table S4 which includes SEMs and *p* values). For WT channels, ΔV_1/2 0→10 mM Mg_ was -60 mV, generally consistent with previous work (7, 18, 27, 50). Proline mutations to the αB helix then reduced the Mg^2+^ induced left shift to 57% - 92% of WT. Hence, an intact αB helix is required for full Mg^2+^ activation.

**Fig. 6.**
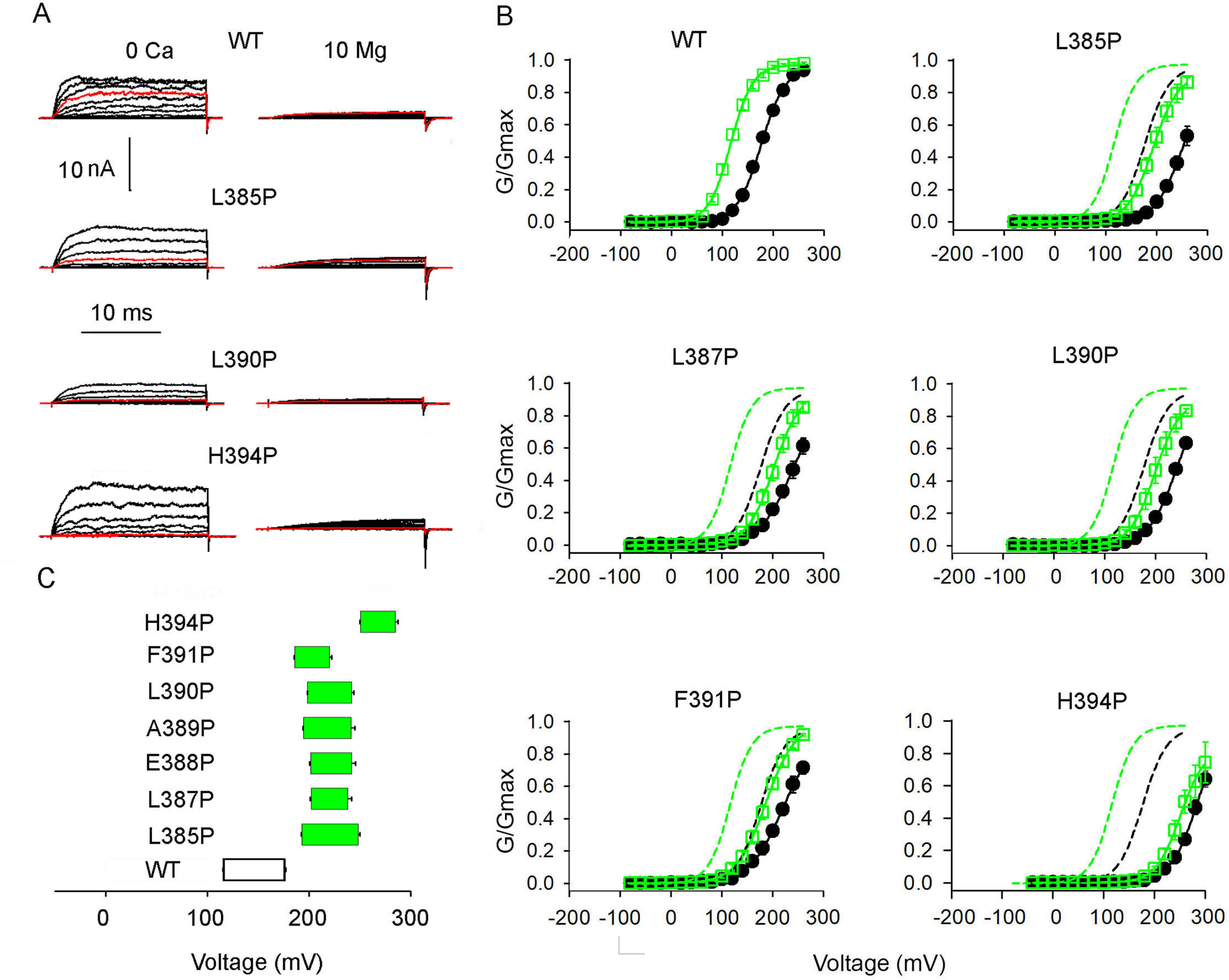
Proline mutations of the αB helix reduce Mg^2+^ activation by an average of 29%. (***A***) Current recordings for the indicated channels. Red traces are currents at 200 mV. Voltage pulses were from -80 to 260 mV, with 20 mV increments, except for H394P where the holding potential was -80 mV and voltage pulses were from -40 to 300 mV. (***B***) G/G_max_ vs. V plots for 0 Mg^2+^ (black lines through black filled circles) and 10 mM Mg^2+^ (green lines through open green squares). Dashed lines are Boltzmann fits for WT channels and continuous lines for mutated channels. Additional data in Fig. S3. Boltzmann fit parameters, SEMs, and *p* values are in *SI Appendix*, Table S4. (***C***) V_1/2_-Mg^2+^ bar plots, where the right end of each bar indicates 0 Mg^2+^ and the left end 10 mM Mg^2+^, as mean and SEM. n≥4.

### Increasing Ca^2+^ 100-fold for Channels with Disrupted αB Helices Does Not Restore High Affinity Ca^2+^ Sensitivity

To explore whether the proline substitutions in the αB helix decreased the Ca^2+^ activation by decreasing the effective occupancy at the high affinity Ca^2+^ binding sites, we increased Ca^2+^ from 100 µM to 10 mM to see if the 100-fold increase in Ca^2+^ would restore the reduced Ca^2+^-induced left shift for αB helix mutated channels to that of WT channels. Results are shown in Fig. 7 and *SI Appendix*, Fig. S4 and Table S2. For αB helix mutated channels, ΔV_1/2 0→100 µM Ca_ is plotted as black bars, and ΔV_1/2 100 µM →10 mM Ca_ is plotted as yellow bars, for comparison to WT channels, plotted as a white bar (Fig. 7*B*). To determine whether the 100-fold increase in Ca^2+^ restores activity at the high affinity Ca^2+^ sites, it is first necessary to know what the 100-fold increase in Ca^2+^ would do at the low affinity Mg^2+^ site. Previous studies have shown that the low affinity Mg^2+^ site is equally activated by 10 mM Ca^2+^ and 10 mM Mg^2+^ (7, 18, 27, 50). Consequently, the effect of 0 to 10 mM Mg^2+^ on delta V_1/2_ (Fig. 6) can be used as a measure of expected effect of 100 µM to 10 mM Ca^2+^ on the low affinity Mg^2+^ site.

**Fig. 7.**
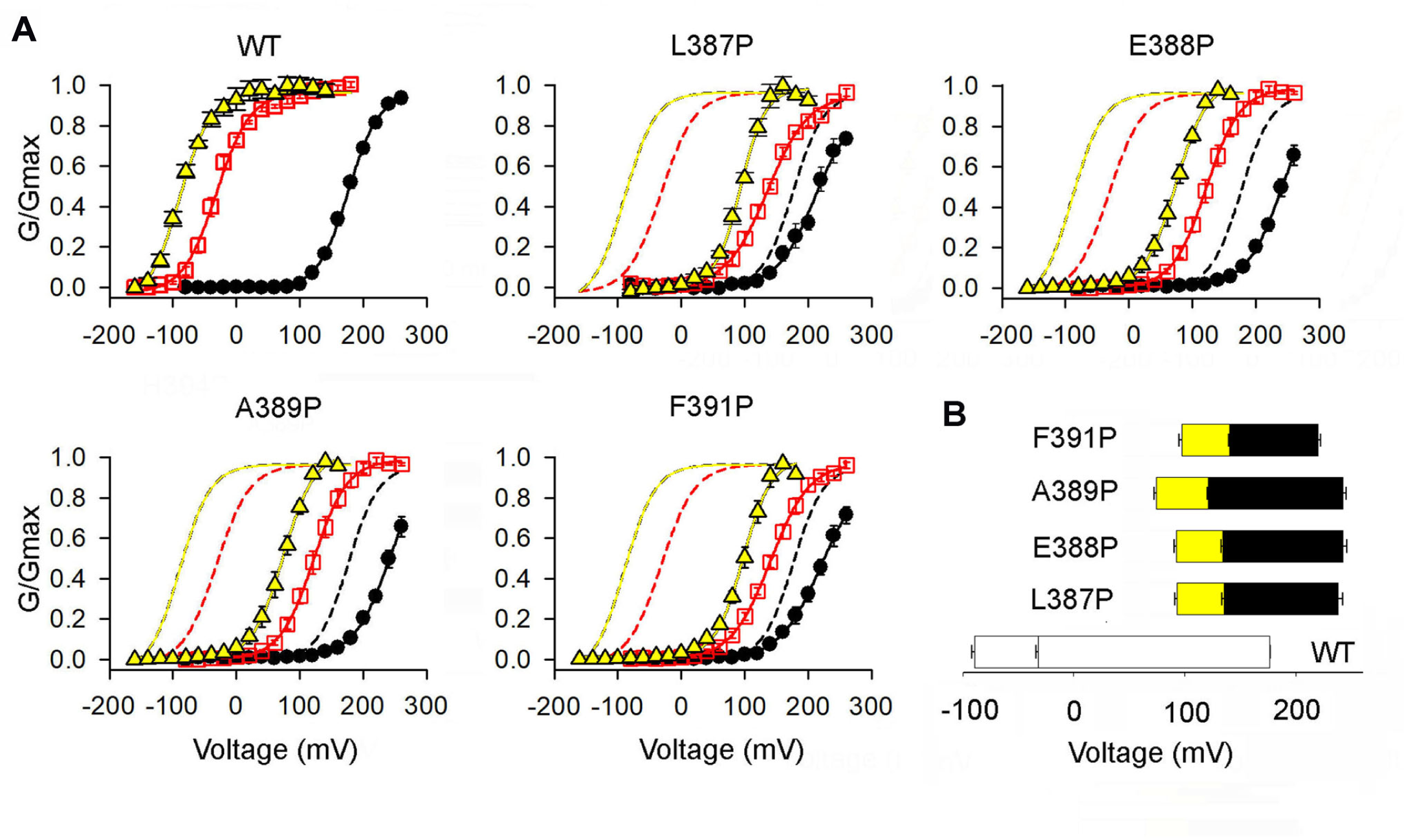
The reduction in Ca^2+^ sensitivity resulting from proline mutations of the αB helix is not reversed by increasing Ca^2+^ 100 fold to 10 mM, suggesting the reduced sensitivity is due to Ca^2+^ activation pathway dysfunction, rather than a reduction in high affinity Ca^2+^ binding. (***A***) G/G_max_ - V plots for the indicated channels in: 0 Ca^2+^ (black lines through filled black symbols); 100 µM Ca^2+^ (red lines through red open squares); and 10 mM Ca^2+^ (yellow lines through yellow filled triangles). Dashed lines are Boltzmann fits to WT channels and continuous lines to mutated channels. Additional data in Fig. S4. Boltzmann fit parameters, SEMs, and *p* values are in *SI Appendix*, Table S2. n≥4. (***B***) V-Ca^2+^ bar plots for V_1/2_ in 0 µM Ca^2+^ (right end of bar), 10 mM Ca^2+^ (left end of bar), and 100 µM Ca^2+^ (line dividing each bar, as mean and SEM).

The ΔV_1/2 100 µM →10 mM Ca_ for the four examined αB helix mutations in Fig. 7 was 46.8 ± 3.2 mV (yellow bars) (data from *SI Appendix*, Table S2). This was not significantly different (*p*=0.14, paired t-test) from ΔV_1/2 0 →10 mM Mg_ of 39.5 ± 3.1 mV for the same four αB helix mutations (Fig. 6, green bars, data from and *SI Appendix*, Table S4). Hence, this comparison suggests that increasing Ca^2+^ 100-fold in Fig. 7 increased activation through the low affinity Mg^2+^ sites with negligible effect on the high affinity Ca^2+^ sites. These observations would be expected if proline mutations to the αB helix decrease Ca^2+^ activation by disrupting a transduction pathway involving the αB helix, rather than by decreasing the affinity of the Ca^2+^ sensors.

## Discussion

### The αB Helix/VSD Interface is Required for Effective Voltage and Ca^2+^ Activation of BK Channels

Functional studies have suggested that voltage and Ca^2+^ sensors of BK channels interact, as summarized in the Introduction (6, 13, 15, 28, 30, 40, 41, 43, 44), and that the AC (N-lobe) region of the CTD is involved (25, 51). Structural studies have identified a potential structural pathway to couple Ca^2+^ sensors in the CTD with voltage sensors in the TMD (16, 17, 31). This pathway includes the interfaces formed between the αB helices at the top of the N-lobes of the CTD and the cytoplasmic sides of the S4-S5 linkers/VSDs (Fig. 1). If this is the case, then disruption of αB helices at the interfaces would be expected to alter both voltage and Ca^2+^ activation (Fig. 1*A, B*).

We explored this possibility by creating an alpha subunit of the tetrameric BK channel that has a proline substitution L390P in the αB helix, with the idea that the proline substitution would disrupt the αB helix, which in turn would alter the interface between the αB helix and S4-S5 linker/VSD. Voltage activation was found to be less for channels formed from the L390P subunits compared to WT channels, as indicated by 66 mV right shifts in the G-V curves. The L390P channels also had a 49% reduction in Ca^2+^ sensitivity, as indicated by a 49% decrease in the left shift of the G-V curves induced by increasing Ca^2+^ from 0 to 100 µM (Fig. 2 and *SI Appendix*, Fig. S1, and Table S2). Both the Ca^2+^ bowl and RCK1 Ca^2+^ site were found to contribute to Ca^2+^sensitivity in WT channels, as previously observed (7, 48). The L390P mutation then reduced the contribution from both of these sites (Fig. 5 and *SI Appendix*, Fig. S2 and Table S3), suggesting that the αB helix may form a common element in an activation pathway used by the two types of high affinity Ca^2+^ sites.

Superposition of crystal structures of WT and mutated CTD showed that the proline mutation L390P near the middle of the αB helix (L390P) caused local disruption of the first turn of the αB helix without significant changes to other parts of the CTD (Fig. 1). Thus, the structural effects of L390P were confined to the αB helix. Single proline mutations at four other examined sites on the αB helix away from the ends, gave results similar to L390P, decreasing both voltage and Ca^2+^ activation (Fig. 2 and *SI Appendix*, Fig. S1 and Table S2). Proline mutations at either end of the αB helix still decreased voltage activation, but had less of an effect on Ca^2+^ activation.

Increasing Ca^2+^ 100 fold to 10 mM did not restore Ca^2+^ sensitivity after proline mutations to the αB helix, indicating that the decrease in Ca^2+^ sensitivity was unlikely to arise from decreased on rates for Ca^2+^ binding at the high affinity Ca^2+^ binding sites, consistent with structural changes from the L390P mutation of the αB helix being local to the αB helix rather than extending to the Ca^2+^ binding sites. Mg^2+^ activation was also decreased by αB helix mutations (Fig. 6 and *SI Appendix*, Fig. S3 and Table S4), but less than for Ca^2+^ activation. The Mg^2+^ coordinating site E399 on the CTD (7, 19, 52) occurs a few residues after the αB helix (Fig. 1*C, D*). Altering the αB helix S4-S5 linker/VSD interface may change the spacing between the Mg^2+^ coordinating sites on the CTD and VSD, which could alter the effectiveness of Mg^2+^ action.

Previous functional observations summarized in the Introduction suggest that voltage and Ca^2+^ sensors interact. Structural observations (16, 17, 31) and our measurements indicating that the αB helices of the Ca^2+^ sensing CTD interface with the S4-S5 linkers/VSDs, together with our observations that a functional αB helix is required for effective voltage and Ca^2+^ activation, suggest that voltage and Ca^2+^ activation may share the αB helix/S4-S5/VSD interface pathway. A common shared element in voltage and Ca^2+^ activation could allow each sensor to share the others’ activation pathway. We now consider potential gating pathways that may contribute to voltage and Ca^2+^ activation of BK channels.

### Potential Voltage Gating Pathways

We first consider two possible pathways for voltage activation: canonical and CTD dependent (17, 30). These pathways are contained in the cartoons in Fig. 1*A, B* and in the Schematic in Fig. 8.

**Fig. 8.**
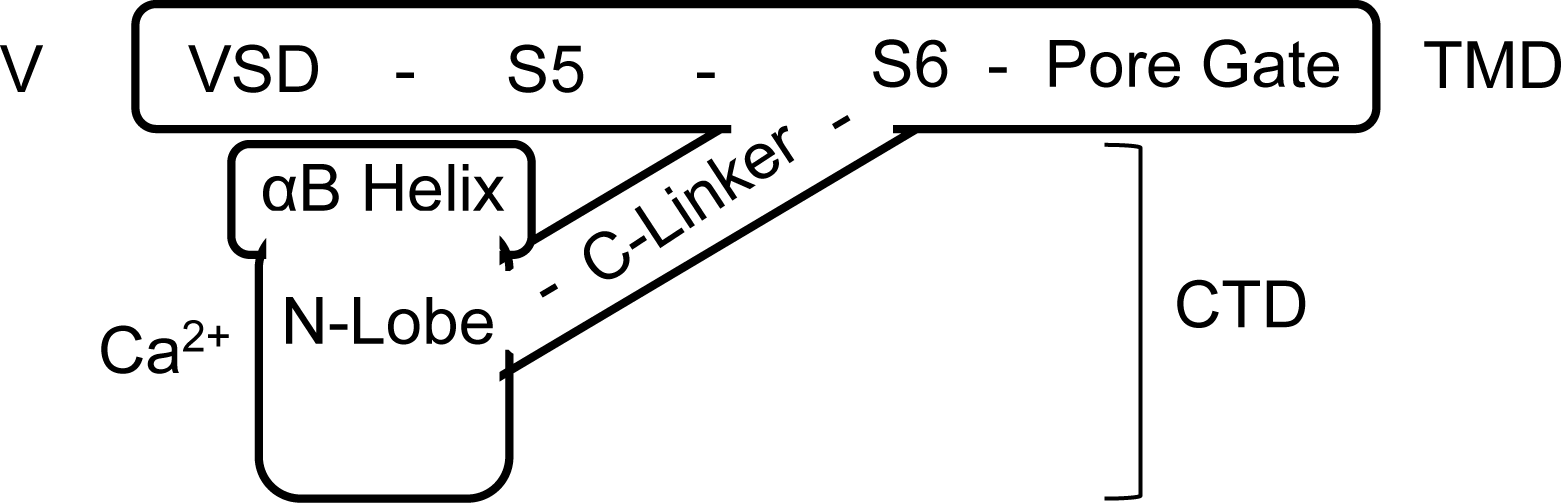
Schematic showing possible voltage and Ca^2+^ transduction pathways. One subunit is shown, as was the case in Fig. 1*A*. In the channel, the αB helix is beneath the S4-S5 linker/VSD of an adjacent subunit in a swapped configuration (16). VSD represents the S4-S5 linker/VSD. Canonical voltage activation would be: V → VSD → S5 → S6 → Pore gate. CTD dependent voltage activation would be: V → VSD → αB helix → N-lobe → C-linker → S6 → Pore gate. C-linker dependent Ca^2+^ activation would be: Ca^2+^ → CTD → N-lobe → C-linker → S6 → pore gate. αB helix dependent Ca^2+^ activation would be: Ca^2+^ → CTD → N-lobe → αB helix → VSD → S5 → S6 → pore gate.

In canonical voltage activation, depolarization elevates S4 in the VSD, which then elevates S5 because of the short S4-S5 linker and extensive interface between S4 and S5 (16, 17). S5 then interacts with S6 to open the pore gate in some manner. This canonical voltage activation pathway is contained within the transmembrane domain: V → VSD → S5 → S6 → pore gate, where VSD includes the S4-S5 linker (Fig. 8). Demonstration of canonical voltage activation in BK channels is the observation that voltage still activates BK channels after removing the CTD (29, 30), which removes the αB helix, N-lobe, and C-linker. However, previous studies showed that voltage activation in the absence of the CTD requires extreme positive voltages (right shifts) for even partial activation of the channel, with Po max from voltage activation being reduced to 29% of WT (29, 30). The Q-V curves were little changed after removing the CTD (left shift of 20 mV), and the coupling factor *D* was decreased 7.2 fold, indicating that the CTD is required to fully couple the voltage sensors to the pore gate for voltage activation (30).

Our current observations provide further support for a role for the CTD in voltage activation. We found that mutating the αB helix in the CTD in channels that were otherwise intact, right shifted the G-V curves (Fig. 2 and *SI Appendix*, Fig. S1 and Table S2), had little effect on the Q-V curves (Fig. 3), and decreased the coupling factor *D* 4.9 fold (Fig. 4), similar to but somewhat less pronounced than the effects of removing the entire CTD. Our observations suggest that the αB helix interface with the S4-S5 linker/VSD is a major contributor to CTD dependent coupling between voltage sensors and the pore gate. We now consider how the CTD might contribute to voltage activation and how disruption of the αB helix could decrease voltage activation through more than one pathway.

In CTD dependent voltage activation, depolarization could elevate S4 of the VSD, altering the relationship of the αB helix with the S4-S5 linker/VSD at their interface (Figs. 1, 8). This alteration of the interface could promote or allow the lateral tilting of the N-lobes away from the central pore. The lateral tilting would move the attachment point of the C-linker on the upper part of the N-lobe laterally and downwards, pulling on S6 to open the pore gate: V → VSD → αB helix → N-lobe → C-linker → S6 → pore gate (Figs, 1, 8), where VSD includes the S4-S5 linker. Disruption of the αB helix would disrupt this CTD depended voltage activation pathway that uses a series C-linker to reach S6 and the pore gate. Support for a CTD dependent voltage activation pathway comes from our observations, discussed above, that mutating the αB helices (Fig. 2 and *SI Appendix*, Fig. S1 and Table S2) or removing the CTD (29, 30) greatly decreases voltage activation through a decrease in the coupling factor *D*.

Further support for a CTD dependent voltage activation pathway comes from the observation that lengthening the C-linker in 0 Ca^2+^ increases the voltage required for activation (39). This is the case because the C-linker is a series component in the CTD dependent voltage activation pathway, being located between the N-lobe and S6 (Fig. 1*A, C* and Fig. 8). As a series component in the CTD dependent activation pathway, lengthening the C-linker would be expected to increase the voltage required for half activation, as more voltage would be required to remove the slack added by lengthening the linker, as observed (39). Whereas this observation supports CTD dependent voltage activation, it is also consistent with the canonical voltage activation pathway. For canonical voltage activation in 0 Ca^2+^, the C-linker-CTD complex would act as a passive spring in parallel with the canonical voltage activation pathway (39). Because the C-linker is very stiff (17), a major component of the spring is likely to arise from the movement of the N-lobe of the CTD (51). Lengthening the C-linker for canonical voltage activation would change the tension in the spring, changing the voltage required for activation, as observed (39). Hence, the observation that increasing C-linker length in 0 Ca^2+^ increases the voltage required for half activation (39) would be consistent with both canonical and CTD dependent voltage activation.

The observations summarized above suggest that both canonical and CTD dependent voltage transduction pathways can contribute to voltage activation. Their relative contributions are likely to depend on Ca^2+^, which moves the N-lobe with the αB helix on top (16, 17), changing the interface area between the αB helix and the S4-S5 linker/VSD (interface areas are presented in the Results). Changing interface area could change the effectiveness of the interface.

### Potential Ca^2+^ Gating Pathways

We next consider two potential pathways for Ca^2+^ activation (16, 17, 31), a C-linker dependent pathway and an αB helix dependent pathway (Fig. 1*A* and Fig. 8). In the C-linker dependent pathway, Ca^2+^ binding to the high affinity Ca^2+^ sites in the CTD could tilt the top of each RCK1 N-lobe laterally away from the central pore. The lateral tilting moves the attachment point of the C-linker on the upper part of each N-lobe laterally and downwards, pulling on S6 to open the pore gate: Ca^2+^ → CTD → N-lobe → C-linker → S6 → pore gate. In the αB helix dependent pathway, the lateral tilting of the N-lobe moves each αB helix upwards and laterally to push upwards and laterally on the VSD and S4-S5 linkers to move S5 and then S6 to open the pore gate: Ca^2+^→ CTD → N-lobe → αB helix → VSD → S5 → S6 → pore gate (Fig. 1*A* and Fig. 8), where the VSD includes the S4-S5 linker. The first three steps in each of these Ca^2+^activation pathways are the same.

Support for Ca^2+^ activation through the C-linker comes from observations that shortening or lengthening the C-linkers increases or decreases the Ca^2+^ activation, respectively (38, 39). Support for the αB helix dependent Ca^2+^ activation pathway comes from our observations that five different proline mutations in the central region of the αB helix decreased Ca^2+^ activation, with ΔV_1/2_ for Ca^2+^ _0 →100 µM_ being decreased to ∼47% of WT (Fig. 2 and *SI Appendix*, Fig. S1 and Table S2). The relative contributions of the C-linker dependent and αB helix dependent Ca^2+^ activation pathways to Ca^2+^ activation are not known. Furthermore, these pathways would interact through an obligatory push-pull mechanism of channel activation (30). The Ca^2+^-induced upwards and lateral movement of the N-lobes would push the αB helices against the bottoms of S4-S5 linkers/VSDs. This push would then push the CTD away from the TMD, increasing the tension in the C-linkers. This increased tension would then further pull on S6 to open the pore gate. Simultaneously, the lateral and downward movement of the C-linker attachment point on the N-lobe would increase the tension in the C-linker, pulling the CTD upwards so that the αB helix would push on the S4-S5 linker/VSD, which could activate the channel through: αB helix → VSD → S5 → S6 → pore gate, where VSD includes the S4-S5 linker. The relative contributions of the two pathways to Ca^2+^ activation, if this is the case, would likely depend on the concentration of Ca^2+^, which increases the interface area between the αB helix and S4-S5 linker/VSD (see Results). Note that a mutation that reduces the push is likely to also reduce pull, and a mutation that reduces the pull is likely to also reduce push (30), which could lead to complications in identifying the relative contributions to Ca^2+^ activation from pathways that change either the push or the pull.

Our observations that the αB helix mutation L390P reduced Ca^2+^ activation for both the Ca^2+^ bowl and the RCK1 Ca^2+^ sites (Fig. 5 and *SI Appendix*, Fig. S2 and Table S3), suggests that both of these high affinity Ca^2+^ sites likely use the N-lobe, αB helix, and S4-S5 linker/VSD as common elements in their activation pathways. Further support for possible common transduction elements comes from the structural observations that Ca^2+^ bowl coordination is completed through contribution of aN438 (hN499) from the RCK1 N-lobe of an adjacent subunit, and that the RCK1 Ca^2+^ site is completed through aD356 (hD367) borrowed from the N-lobe of the same subunit, such that the two high affinity sites may work jointly to tilt the N-lobe laterally and upward as a common transduction element for the RCK1 and Ca^2+^ bowl sites (16, 17). Functional studies further indicate that hR514 from RCK1 may coordinate with Ca^2+^ bowl hE902 and hY904, leading to additional potential common elements in the Ca^2+^ bowl and RCK1 Ca^2+^ site activation pathways (48).

However, there are also indications of some potential differences in activation pathways, as Ca^2+^ binding is voltage dependent at the RCK1 Ca^2+^ site, but not at the Ca^2+^ bowl site (41), and strong correlation has been observed between VSD function and RCK1 conformational changes, but not between RCK2 and VSD function (44). Additional evidence for some possible differences in the RCK1 Ca^2+^ site activation pathway and the Ca^2+^ bowl activation pathway are our observations that disruption of the αB helix with L390P reduced the fractional contribution to Ca^2+^ sensitivity from RCK1 Ca^2+^ sites more than from Ca^2+^ bowl sites (Fig. 5 and *SI Appendix*, Fig. S2 and Table S3). Such a differential reduction could occur if the RCK1 Ca^2+^ site utalized the αB helix dependent Ca^2+^ activation pathway more than the C-linker Ca^2+^ dependent pathway.

### αB Helix Mutations Decrease Mg^2+^ Activation of BK channels

Mg^2+^ is coordinated between E374 and E399 on top of the N-lobe and N172 and other residues on the VSD, linking Ca^2+^ sensors in the CTD with voltage sensors (16, 40, 53). Ten mM Mg^2+^ activated BK channels with a -60 mV shift in the G-V curves (Fig. 6 and *SI Appendix*, Fig. S3 and Table S4). We found that seven different proline mutations in the αB helix reduced Mg^2+^ activation to 57% - 92% of WT, with L390P reducing activation to 72% of WT (Fig. 6 and *SI Appendix*, Fig. S3 and Table S4), without significant changes in structure of the RCK1-RCK2 domains away from the αB helix (Fig. 1*E, F*). These observations suggest that an intact αB helix S4-S5 linker/VSD interface is required for effective Mg^2+^ activation. If Mg^2+^ binding electrostatically pulls the CTD with attached αB helix closer to the S4-S5 linker/VSD, this could increase the push of the αB helix on the S4-S5 linker/VSD, potentially increasing activation. Mutating the αB helix could then decrease this activation, as observed.

### Conclusion

Our experiments show that an interface formed between the αB helices at the top of the cytosolic CTD and the bottom of the transmembrane voltage sensor domains plays a critical role in both voltage and Ca^2+^ activation of BK channels. This αB helix/voltage sensor domain interface could couple voltage and Ca^2+^ transduction pathways, allowing voltage and Ca^2+^ to activate through both independent pathways, and pathways with some shared components.

## Materials and Methods

### Structure determination of human BK CTD L390P mutant

We determined the crystal structure of the L390P mutant (PDB:6V5A) by molecular replacement using the human BK CTD (PDB: 3MT5) with a deletion of the αB helix (residues 384-394) as a search model. The αB helix harboring the L390P mutation was built near the end of structural refinement to reduce model bias from molecular replacement. The final model was refined to a resolution of 2.0 Å with R_work_/R_free_= 20.3%/22.5%.

### Electrophysiology recording

Site-directed mutagenesis and PCR were used to make the constructs from the mbr5 splice of variant of mSlo1 (3). cRNA was synthesized and injected into Xenopus laevis oocytes for channel expression. Inside-out patch clamp recording (30) was used to record ionic currents and gating currents, which were digitized and analyzed with pClamp and HEKA Elektronik systems. Solver in Excel was used for fitting the HCA model. For ionic currents the pipette (external) solution contained (mM): 150 KCl, 2 MgCl_2,_ 5 TES buffer. The 0 Ca^2+^ internal solution contained (mM): 150 KCl, 1 EGTA, 1 HEDTA, and 5 TES, pH 7.0. The 100 µM internal Ca^2+^ solution contained (mM), 150 KC1, 0.1 CaCl_2_, 5 TES, pH 7.0. The 10 mM internal Ca^2+^ solution contained (mM): 150 KC1, 10 CaCl_2_, 5 TES, pH 7.0. The 10 mM CaCl_2_ was replaced by 10 mM MgCl_2_ for the 10 mM internal Mg^2+^ solution. Experiments were at 22°C to 24°C. Error bars are SEM.

**Additional information on materials and methods is provided in *SI Appendix***

## ACKNOWLEDGEMENTS

This work was support by NIH grants R01-GM114694 to L.S. and K.L.M. and R01-HL142301 to J.C.

## References

1. J. N. Barrett, K. L. Magleby, B. S. Pallotta, Properties of single calcium-activated potassium channels in cultured rat muscle. J. Physiol 331, 211–230 (1982).

2. E. Moczydlowski, R. Latorre, Gating kinetics of Ca^2+^-activated K^+^ channels from rat muscle incorporated into planar lipid bilayers. Evidence for two voltage-dependent Ca^2+^ binding reactions. J. Gen. Physiol 82, 511–542 (1983).

3. A. Butler, S. Tsunoda, D. P. McCobb, A. Wei, L. Salkoff, mSlo, a complex mouse gene encoding “maxi” calcium-activated potassium channels. Science 261, 221–224 (1993).

4. M. Schreiber, L. Salkoff, A novel calcium-sensing domain in the BK channel. Biophys. J 73, 1355–1363 (1997).

5. B. S. Rothberg, K. L. Magleby, Voltage and Ca^2+^ activation of single large-conductance Ca^2+^-activated K^+^ channels described by a two-tiered allosteric gating mechanism. J. Gen. Physiol 116, 75–99 (2000).

6. F. T. Horrigan, R. W. Aldrich, Coupling between voltage sensor activation, Ca^2+^ binding and channel opening in large conductance (BK) potassium channels. J. Gen. Physiol 120, 267–305 (2002).

7. X. M. Xia, X. Zeng, C. J. Lingle, Multiple regulatory sites in large-conductance calcium-activated potassium channels. Nature 418, 880–884 (2002).

8. L. Bao, A. M. Rapin, E. C. Holmstrand, D. H. Cox, Elimination of the BK(Ca) channel’s high-affinity Ca^2+^ sensitivity. J. Gen. Physiol 120, 173–189 (2002).

9. U. S. Lee, J. Shi, J. Cui, Modulation of BK channel gating by the ss2 subunit involves both membrane-spanning and cytoplasmic domains of Slo1. J. Neurosci 30, 16170–16179 (2010).

10. T. Hoshi, A. Pantazis, R. Olcese, Transduction of voltage and Ca^2+^ signals by Slo1 BK channels. Physiology. (Bethesda.) 28, 172–189 (2013).

11. Y. Geng, K. L. Magleby, Single-channel kinetics of BK (Slo1) channels. Front Physiol 5, 532 (2014).

12. H. Yang, G. Zhang, J. Cui, BK channels: multiple sensors, one activation gate. Front Physiol 6, 29 (2015).

13. A. Pantazis, R. Olcese, Biophysics of BK Channel Gating. Int. Rev. Neurobiol 128, 1–49 (2016).

14. R. Latorre et al., Molecular Determinants of BK Channel Functional Diversity and Functioning. Physiol Rev 97, 39–87 (2017).

15. Y. Zhou, H. Yang, J. Cui, C. J. Lingle, Threading the biophysics of mammalian Slo1 channels onto structures of an invertebrate Slo1 channel. J. Gen. Physiol 149, 985–1007 (2017).

16. X. Tao, R. K. Hite, R. MacKinnon, Cryo-EM structure of the open high-conductance Ca^2+^-activated K^+^ channel. Nature 541, 46–51 (2017).

17. R. K. Hite, X. Tao, R. MacKinnon, Structural basis for gating the high-conductance Ca^2+^-activated K^+^ channel. Nature 541, 52–57 (2017).

18. X. Zhang, C. R. Solaro, C. J. Lingle, Allosteric regulation of BK channel gating by Ca^2+^ and Mg^2+^ through a nonselective, low affinity divalent cation site. J. Gen. Physiol 118, 607–636 (2001).

19. J. Yang et al., Interaction between residues in the Mg^2+^-binding site regulates BK channel activation. J. Gen. Physiol 141, 217–228 (2013).

20. R. Brenner et al., Vasoregulation by the beta1 subunit of the calcium-activated potassium channel. Nature 407, 870–876 (2000).

21. R. Robitaille, M. L. Garcia, G. J. Kaczorowski, M. P. Charlton, Functional colocalization of calcium and calcium-gated potassium channels in control of transmitter release. Neuron 11, 645–655 (1993).

22. N. Gu, K. Vervaeke, J. F. Storm, BK potassium channels facilitate high-frequency firing and cause early spike frequency adaptation in rat CA1 hippocampal pyramidal cells. J Physiol 580, 859–882 (2007).

23. J. D. Holtzclaw, P. R. Grimm, S. C. Sansom, Role of BK channels in hypertension and potassium secretion. Current Opinion in Nephrology and Hypertension 20, 512–517 (2011).

24. B. Wang, B. S. Rothberg, R. Brenner, Mechanism of increased BK channel activation from a channel mutation that causes epilepsy. J. Gen. Physiol 133, 283–294 (2009).

25. J. Yang et al., An epilepsy/dyskinesia-associated mutation enhances BK channel activation by potentiating Ca^2+^ sensing. Neuron 66, 871–883 (2010).

26. H. Jiao et al., Genome wide association study identifies KCNMA1 contributing to human obesity. Bmc Medical Genomics 4 (2011).

27. J. Shi, J. Cui, Intracellular Mg^2+^ enhances the function of BK-type Ca^2+^-activated K^+^ channels. J. Gen. Physiol 118, 589–606 (2001).

28. H. Yang et al., Activation of Slo1 BK channels by Mg^2+^ coordinated between the voltage sensor and RCK1 domains. Nat. Struct. Mol. Biol 15, 1152–1159 (2008).

29. G. Budelli, Y. Geng, A. Butler, K. L. Magleby, L. Salkoff, Properties of Slo1 K^+^ channels with and without the gating ring. Proc. Natl. Acad. Sci. U. S. A 110, 16657–16662 (2013).

30. G. Zhang et al., Deletion of cytosolic gating ring decreases gate and voltage sensor coupling in BK channels. J. Gen. Physiol 149, 373–387 (2017).

31. P. Yuan, M. D. Leonetti, Y. C. Hsiung, R. MacKinnon, Open structure of the Ca^2+^ gating ring in the high-conductance Ca^2+^-activated K^+^ channel. Nature 481, 94–U105 (2012).

32. O. M. Koval, Y. Fan, B. S. Rothberg, A role for the S0 transmembrane segment in voltage-dependent gating of BK channels. J. Gen. Physiol 129, 209–220 (2007).

33. Z. Jia, M. Yazdani, G. Zhang, J. Cui, J. Chen, Hydrophobic gating in BK channels. Nat. Commun 9, 3408 (2018).

34. X. Chen, J. Yan, R. W. Aldrich, BK channel opening involves side-chain reorientation of multiple deep-pore residues. Proc. Natl. Acad. Sci. U. S. A 111, E79–E88 (2014).

35. Y. Wu, Y. Yang, S. Ye, Y. Jiang, Structure of the gating ring from the human large-conductance Ca^2+^-gated K^+^ channel. Nature 466, 393–397 (2010).

36. X. H. Zeng, X. M. Xia, C. J. Lingle, Divalent cation sensitivity of BK channel activation supports the existence of three distinct binding sites. J. Gen. Physiol 125, 273–286 (2005).

37. Z. Ma, X. J. Lou, F. T. Horrigan, Role of charged residues in the S1-S4 voltage sensor of BK channels. J. Gen. Physiol 127, 309–328 (2006).

38. A. R. Pico, RCK Domain Model of Calcium Activation in BK Channels (Rockefeller University, New York, 2003).

39. X. Niu, X. Qian, K. L. Magleby, Linker-gating ring complex as passive spring and Ca^2+^-dependent machine for a voltage- and Ca^2+^-activated potassium channel. Neuron 42, 745–756 (2004).

40. U. S. Lee, J. Cui, BK channel activation: structural and functional insights. Trends Neurosci 33, 415–423 (2010).

41. T. B. Sweet, D. H. Cox, Measurements of the BKCa channel’s high-affinity Ca^2+^ binding constants: effects of membrane voltage. J. Gen. Physiol 132, 491–505 (2008).

42. N. Savalli, A. Pantazis, T. Yusifov, D. Sigg, R. Olcese, The Contribution of RCK Domains to Human BK Channel Allosteric Activation. Journal of Biological Chemistry 287, 21741–21750 (2012).

43. Y. Lorenzo-Ceballos, W. Carrasquel-Ursulaez, K. Castillo, O. Alvarez, R. Latorre, Calcium-driven regulation of voltage-sensing domains in BK channels. Elife 8 (2019).

44. P. Miranda, M. Holmgren, T. Giraldez, Voltage-dependent dynamics of the BK channel cytosolic gating ring are coupled to the membrane-embedded voltage sensor. Elife 7 (2018).

45. F. T. Horrigan, J. Cui, R. W. Aldrich, Allosteric voltage gating of potassium channels I. Mslo ionic currents in the absence of Ca^2+^. J. Gen. Physiol 114, 277–304 (1999).

46. J. Cui, D. H. Cox, R. W. Aldrich, Intrinsic voltage dependence and Ca^2+^ regulation of mslo large conductance Ca-activated K^+^ channels. J. Gen. Physiol 109, 647–673 (1997).

47. R. Guan et al., Allosteric-activation mechanism of BK channel gating ring triggered by calcium ions. PLoS One 12, e0182067 (2017).

48. A. S. Kshatri, A. J. Gonzalez-Hernandez, T. Giraldez, Functional validation of Ca^2+^-binding residues from the crystal structure of the BK ion channel. Biochim. Biophys. Acta Biomembr 1860, 943–952 (2018).

49. Q. Li et al., Molecular determinants of Ca^2+^ sensitivity at the intersubunit interface of the BK channel gating ring. Sci. Rep 8, 509 (2018).

50. L. Hu, H. Yang, J. Shi, J. Cui, Effects of multiple metal binding sites on calcium and magnesium-dependent activation of BK channels. J. Gen. Physiol 127, 35–49 (2006).

51. G. Krishnamoorthy, J. Shi, D. Sept, J. Cui, The NH2 terminus of RCK1 domain regulates Ca^2+^-dependent BK(Ca) channel gating. J. Gen. Physiol 126, 227–241 (2005).

52. J. Shi et al., Mechanism of magnesium activation of calcium-activated potassium channels. Nature 418, 876–880 (2002).

53. H. Yang et al., Mg^2+^ mediates interaction between the voltage sensor and cytosolic domain to activate BK channels. Proc. Natl. Acad. Sci. U. S. A 104, 18270–18275 (2007).

54. P. Yuan, M. D. Leonetti, A. R. Pico, Y. Hsiung, R. MacKinnon, Structure of the human BK channel Ca^2+^-activation apparatus at 3.0 A resolution. Science 329, 182–186 (2010).

