## Supplementary material for "Coupling of Ca^2+^ and voltage activation in BK channels through the αB helix/voltage sensor interface": SI Appendix

**This PDF file includes:**

Supplementary Materials and Methods
Figures S1 to S4
Tables S1 to S4
SI References

### Supporting Information

### Materials and Methods

**Structural Determination of the Human BK CTD L390P Mutant.** The DNA encoding the human BK CTD (1) was ligated into the EcoR1/Xho1 restriction sites of a modified pPICZ-B vector (Invitrogen). The resulting protein including residues 341-1056 was fused to a C-terminal GFP-$His_{10}$ tag linked by a PreScission cleavage site. The L390P mutant construct was generated by site-directed mutagenesis and the protein was expressed in *Pichia pastoris*. Cells were first disrupted by milling and then were re-suspended in lysis buffer containing 50 mM Tris pH 8.0, 300 mM NaCl and 5 mM 2-mecaptoethanol supplemented with protease inhibitors including 2.5 µg/mL Leupeptin, 1 µg/mL Pepstatin A, 100 µg/mL 4-(2-Aminoethyl) benzenesulfonyl fluoride hydrochloride, 3 µg/mL Aprotinin, 1 mM Benzamidine and 200 µM phenylmethane sulphonylfluoride. The lysate was further disrupted by sonication and then was centrifuged for 1 hour at 30,000 *g*. Supernatant was loaded onto TALON resin and washed with buffer containing 10 mM imidazole and then eluted with 300 mM imidazole. The eluted protein was incubated overnight with PreScission protease to remove the C-terminal GFP-$His_{10}$ tag and was further purified using a Superose 6 column (GE Healthcare) pre-equilibrated with buffer containing 20 mM Tris pH 8.0, 150 mM NaCl, 5 mM dithiothreitol (DTT), and 50 mM $CaCl_2$. Peak fractions were collected and concentrated to ~ 6.5 mg/ml for crystallization experiments.

Crystals of the L390P mutant were grown at 22°C using hanging-drop vapor diffusion by mixing 0.3 µL protein with 0.3 µL reservoir solution containing 50 mM sodium acetate, 4% (w/v) PEG 4000, 100 mM sodium sulfate, and 100 mM lithium sulfate (pH 4.9). All crystals were cryo-protected in the reservoir solution supplemented with 150 mM NaCl, 50 mM $CaCl_2$ and 30% ethylene glycol, and flashed-cooled in liquid nitrogen.

Diffraction data were collected at the Advanced Photon Source beamline 24-ID-C and were processed with the HKL2000 program suite (2). A molecular replacement solution was obtained using Phaser (3) with the human BK CTD (PDB: 3MT5) lacking the αB helix (residues 384-394) as a search model. Cycles of model refinement were carried out in Coot (4) and REFMAC (5) before modeling the missing αB helix to reduce model bias from molecular replacement. The αB helix harboring the L390P mutation was built near the end of structural refinement. The final model was refined to a resolution of 2.0 Å with $R_{work}/R_{free}$= 20.3%/22.5%. The overall structure of the L390P mutant is very similar to that of the wild type, with an RMSD of 0.6 Å for all Cα atoms. In the L390P structure, introduction of a proline residue in the middle of the αB helix results in a shorter αB helix. The remaining structure, including the $Ca^{2+}$-bowl, is nearly identical to that of the wild type.

Protein interaction interfaces (buried surface area) were measured between the αB helix and the S4-S5 linker and the VSD for both closed (PDB 5TJI, EDTA) and open (PDB 5TJ6, $Ca^{2+}$ and $Mg^{2+}$ liganded) structures of Aplysia Slo1 channels (6, 7). The αB helilx, S4-S5 linker, and combined S4-S5 linker/VSD were defined as Aplysia residues 373-383, 214-219, and 14-219, respectively. Surface area estimations were done using AREAIMOL in CCP4 (8).

**Expression Construct and Molecular Biology.** Site-directed mutagenesis using Stratagene's Quick Change Mutagenesis Kit, and overlap extension PCR were used to make the constructs from the mbr5 splice variant of mSlo1 (9). cRNA was synthesized with T3 polymerase in vitro (Ambion). 1ng to 50ng/cell RNA was injected into Xenopus laevis oocytes, and the injected oocytes were incubated in the Barth's solution at 18°C for 2 to 5 days before currents were recorded. Barth's solution (in mM): 88 NaCl, 1 KCl, 0.4 $CaCl_2$, 0.33 $Ca(NO_3)_2$, 0.8 $MgSO_4$, 2.4 $NaHCO_3$, 5 TES, pH7.2.

**Gating Current Recording.** Gating currents were recorded from inside-out patches (10). The gating currents were sampled at 200 kHz and filtered at 20 kHz, with leak subtraction using a −P/4 protocol. The pipette solution contained (mM): 125 methanesulfonic acid, 127 TEA hydroxide, 2 HCl, 2 $MgCl_2$, and 20 HEPES. The internal solution contained (mM): 135 methanesulfonic acid, 141 NMDG, 6 HCl, 20 HEPES, and 5 EGTA, pH 7.2. Stage V or VI oocytes were obtained from Xenopus laevis by laparotomy. All procedures were performed in accordance with the protocol approved by the Washington University Animal Studies Committee.

**Ionic Current Recording.** Inside-out patches were used to record ionic currents with an Axopatch 200-B patch-clamp amplifier (Molecular Devices) and pClamp 9.0 software. Borosilicate pipettes with 0.5–2 MΩ resistance were used. The current signals were filtered at 10 KHz and were sampled at 100 kHz. A -P/4 protocol was used to remove capacitive transients and leak currents. All experiments were done at room temperature of 22°C to 24 °C. The pipette (external) solution contained (mM): 150 KCl, 2 $MgCl_2$, 5 TES buffer. The 0 $Ca^{2+}$ internal solution contained (mM): 150 KCl, 1 EGTA, 1 HEDTA, and 5 TES, pH 7.0. The 100 μM internal $Ca^{2+}$ solution contained (mM), 150 KC1, 0.1 $CaCl_2$, 5 TES, pH 7.0. The 10 mM internal $Ca^{2+}$ solution contained (mM): 150 KC1, 10 $CaCl_2$, 5 TES, pH 7.0. The 10 mM $CaCl_2$ was replaced with 10 mM $MgCl_2$ for the 10 mM internal $Mg^{2+}$ solution. Procedures to obtain oocytes from Xenopus laevis were approved by the University of Miami Animal Care and Use Committee.

**Data Analysis for G-V Curves.** Relative conductance was determined from macroscopic tail current amplitudes using the voltage protocols indicated in the figure legends. $G/G_{max}$ vs. V curves (G-V) were fitted with the Boltzmann function (10) to estimate $V_{1/2}$, the voltage required for half maximal activation, and *b*, the voltage dependence of activation, using

$$G/G_{max} = 1/(1 + \exp( (V_{1/2} - V)\, z e_o / k_B T)) \quad (1)$$

$$G/G_{max} = 1/(1 + \exp(V_{1/2} - V)/b) \quad (2)$$

where $G/G_{max}$ is the ratio of conductance to maximum conductance, *z* is the number of equivalent units of elementary charge associated with gating, V is membrane potential, k is Boltzmann's constant, T is absolute temperature, *b* is the slope factor, which indicates the chance in millivolts required to increase open probability (Po) *e*-fold at very low Po, $k_BT/e_o$ has a value of 25.52 mV at 23°C, and *z* is given by 25.52 mV/*b*. It should be noted that estimates of *z* obtained from fitting the slopes of G-V curves are typically underestimated by a factor of two compared to estimates obtained from fitting Po-V data from single channels (11). This underestimation arises from the variability in $V_{1/2}$ among the large number of channels averaged by macro-patch recordings, which decreases the slope of the G-V curves. All experiments were repeated four or more times. Plotted data points in G-V and Q-V plots are mean ± SEM of data averaged from different experiments. Error estimates of the SEM for $V_{1/2}$ and *z* where from the Boltzmann fits to the average data points for G-V and Q-V plots.

$Ca^{2+}$ sensitivity was measured by the magnitude of the left shift in $V_{1/2}$ induced by increasing $Ca^{2+}$ from 0 to 100 μM (12), such that

$$\Delta V_{1/2\ 0\rightarrow 100\ Ca} = V_{1/2\ 100\ Ca} - V_{1/2\ 0\ Ca} \quad (3)$$

where $V_{1/2\ 0\ Ca}$ and $V_{1/2\ 100\ Ca}$ are the voltages for half maximal activation determined with 0 and 100 μM $Ca^{2+}$. Low affinity $Ca^{2+}$ activation was determined by replacing 0 $Ca^{2+}$ and 100 μM $Ca^{2+}$ in Eqn. 3 with 100 μM and 10 mM $Ca^{2+}$. Low affinity $Mg^{2+}$ activation was determined by replacing 0 and 100 μM $Ca^{2+}$ in Eqn. 3 with 0 and 10 mM $Mg^{2+}$.

Data points in the $G/G_{max}$ vs. voltage plots (G-V plots) are plotted as mean ± SEM for the values determined in separate experiments. $V_{1/2}$ and *z* and their error bars as SEM were determined from the Boltzmann fits to the average data points in the G-V plots using SigmaPlot 14. $\Delta V_{1/2}$ ± SEM values were determined using GraphPad Prism8.1.1. Two tailed *t* tests were performed using the Holm-Sidak method which corrects the *p* values for multiple comparisons.

**Po versus V Plots.** To obtain the Po versus V plots over wide ranges of Po, patches containing hundreds to thousands of channels were stimulated with a 5 s step at each voltage. A baseline was subtracted from the recorded current records (examples in Fig. 4*B*) and all open events were then integrated to obtain an area of current times open time. This was then divided by single-channel current amplitude and total time to obtain NPo, the average Po for all channels in the patch, as described previously (10, 13), where N, the total number of channels in the patch, was estimated from the macroscopic current at 250 mV at 0 μM $Ca^{2+}$.

**Estimating *D* and $L_0$.** The HCA model (13) for the steady-state voltage dependent gating of BK channels (Fig. 4*A*) was used to estimate *D*, the allosteric factor that couples voltage sensor activation to channel opening and $L_0$, the intrinisic closed to open equilibrium constant at 0 mV. For the HCA model open probability Po is given by (13)

$$Po = 1/(1+(1+J)^4/L(1+DJ)^4) \quad (4)$$

where *J* is the equilibrium constant for voltage sensor activation in each subunit, and $L_0$ is the intrinsic equilibrium constant for closed to open transitions. Both *J* and *L* are voltage dependent, such that

$$J = \exp((V-V_{hc})Z_J/k_BT), \text{ and} \quad (5)$$

$$L = L_0\exp(Z_LV/k_BT) \quad (6)$$

where $L_0$ is the value of $L$ at 0 mV, $Z_J$ and $Z_L$ are the effective charge movement for equilibrium constants $J$ and $L$, and $V$hc is the voltage for half-activation of the on-gating current which gives the half-activation voltage at the closed state. The gating current versus voltage plot gives $V_{hc}$ and $Z_J$ from the Boltzman fits.

The mean Po-V data in Fig. 4*C* was fitted using Solver in Excel (10) with simultaneous Eqs. 4, 5 and 6 with fixed parameters $V_{hc}$ and $Z_J$ to determine $D$, $L_0$, and $Z_L$. Estimates of SEM for these parameters were determined by individually fitting data from four separate experiments.

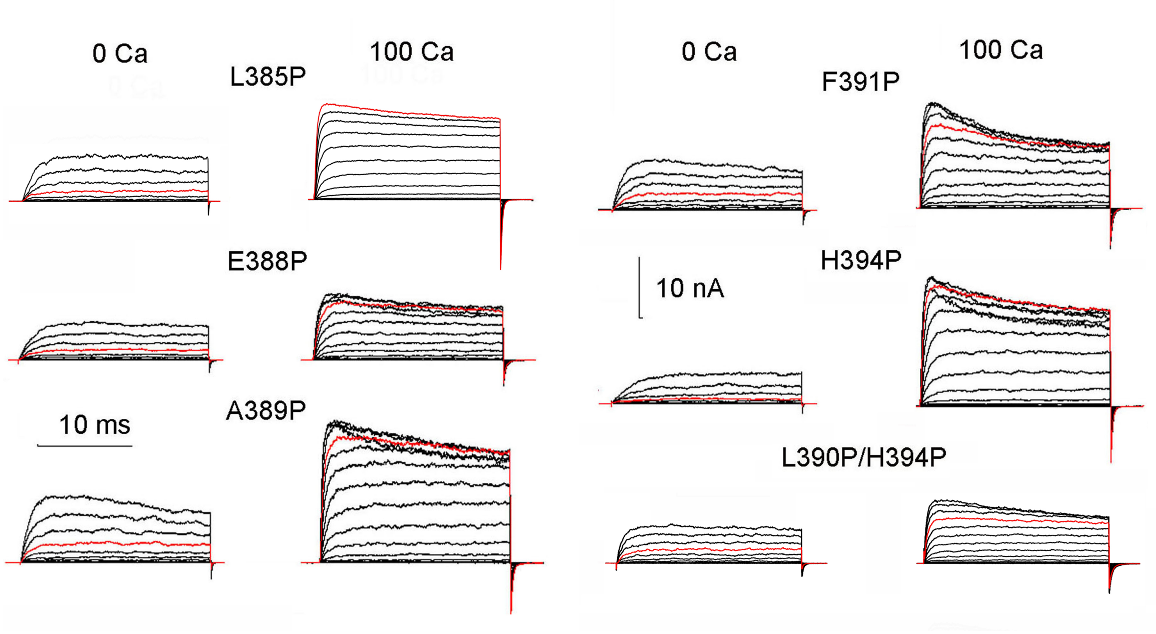


**Fig. S1.** Supplemental data for Fig. 2. Proline mutations of the αB helix decrease both voltage and $Ca^{2+}$ activation. Current recordings for the indicated channels from inside-out macro patches. Red traces are currents at 200 mV. Voltage pulses were from -80 to 260 mV with a 20 mV increments, except for L385P with 100 μM $Ca^{2+}$ which were from -140 mV to 200 mV.

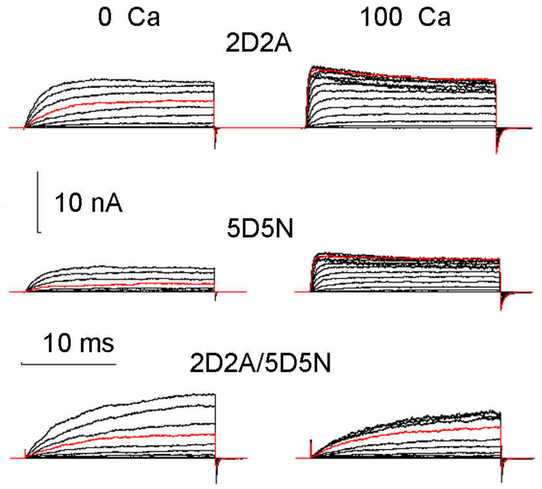


**Fig. S2.** Supplemental data for Fig. 5. Both the $Ca^{2+}$ bowl and RCK1 $Ca^{2+}$ binding sites require an intact αB helix for effective $Ca^{2+}$ activation. Current recordings from the indicated channels. Red traces are currents at 200 mV. Voltage pulses were from -80 to 260 mV with 20 mV increments.

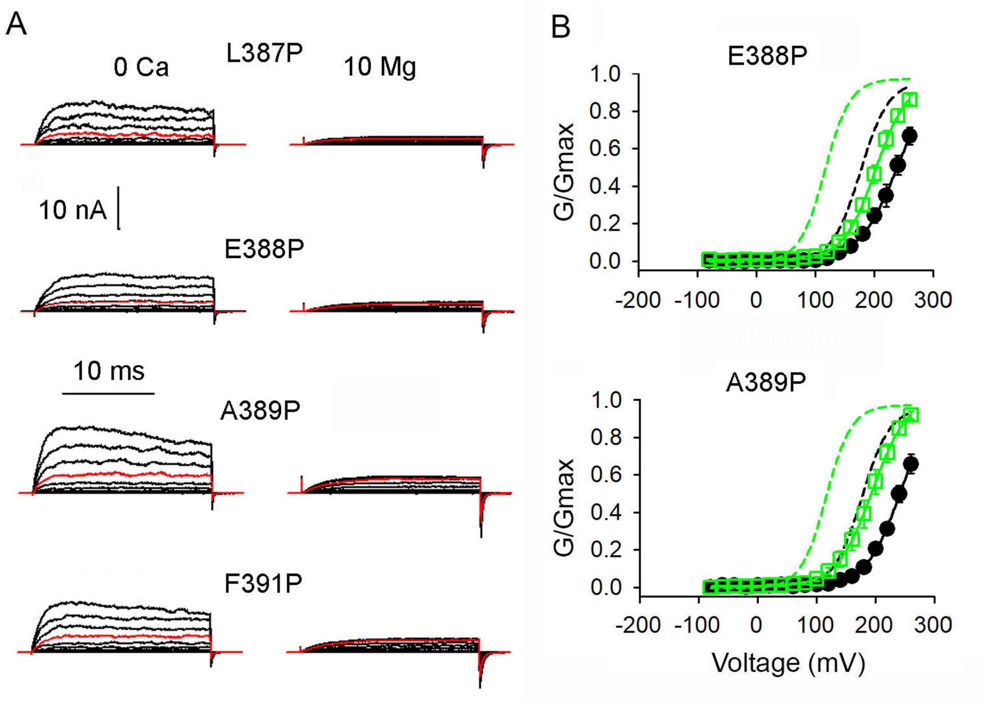


**Fig. S3.** Supplemental data for Fig. 6. Proline mutations of the αB helix reduce $Mg^{2+}$ activation by an average of 29%. (A) Current recordings from the indicated channels. Red traces are currents at 200 mV. Voltage pulses were from -80 to 260 mV with 20 mV increments. (B) G/Gmax vs. V plots in 0 $Ca^{2+}$ (black lines through filled black circles, n≥4) and 10 mM $Mg^{2+}$ (green lines through open squares, n≥4). Dashed lines are Boltzmann fits for WT channels and continuous lines are for mutated channels. Boltzman parameters in Table S3.

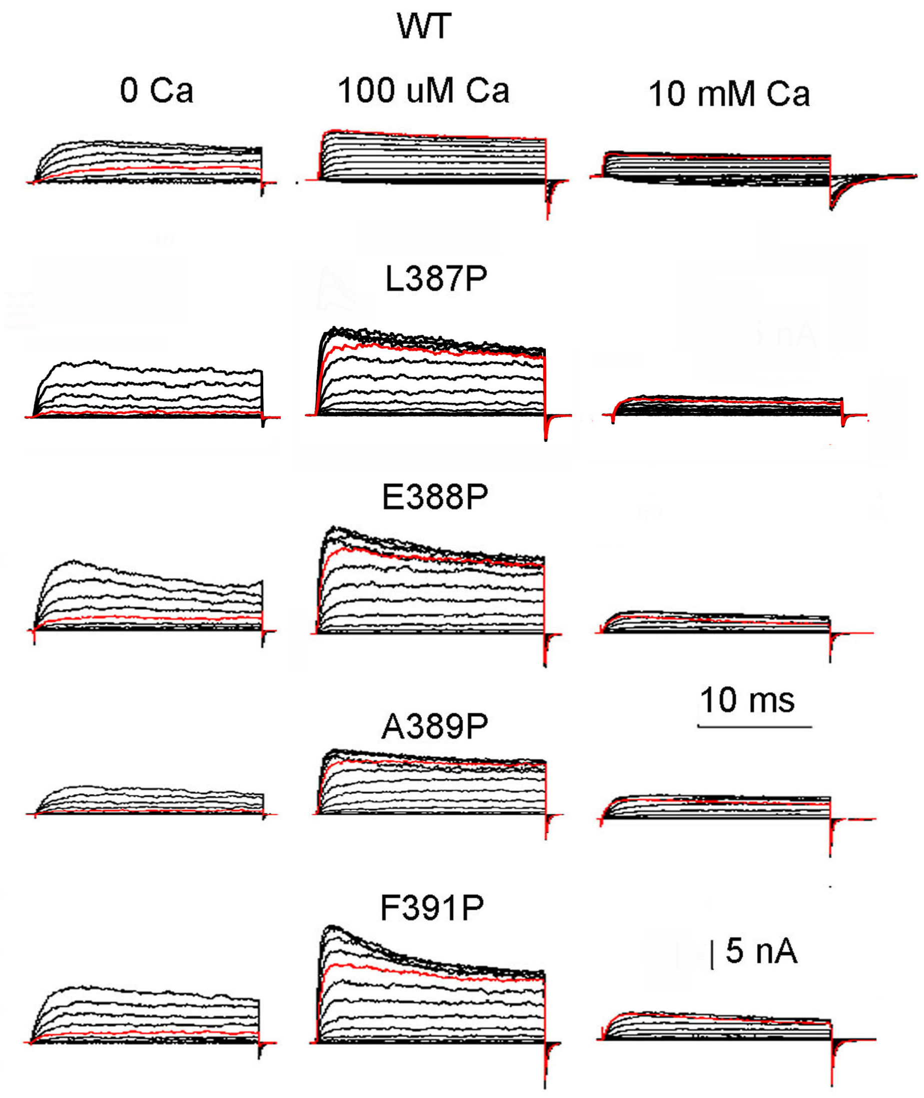


**Fig. S4.** Supplemental data for Fig. 7. The reduction in $Ca^{2+}$ sensitivity resulting from proline mutations of the αB helix is not reversed by increasing $Ca^{2+}$ 100 fold. Current recordings for the indicated channels. Red traces are currents at 180 mV. Voltages pulses were from -80 to 260 mV with 20 mV increments, except for WT channels at 100 μM and 10 mM $Ca^{2+}$ where the voltage pulses were from -160 to 180 mV. The scale bar for L387P is 1.5 nA.

**Table S1. Summary of crystallographic data**

| **Crystal** | hBK CTD L390P |
|---|---|
| **Data collection** | |
| Space group | H32 |
| Cell dimensions | |
| *a*, *b*, *c* (Å) | 145.5, 145.5, 242.9 |
| α, β, γ (˚) | 90, 90, 120 |
| Resolution (Å) | 2.0 |
| $R_{merge}$ | 0.055 (0.734) |
| $I/\sigma I$ | 30.6 (2.4) |
| Completeness (%) | 100.0 (100.0) |
| Redundancy | 8.2 (7.5) |
| **Refinement** | |
| Resolution (Å) | 30 - 2.0 |
| No. of reflections (total/test) | 63458/3340 |
| $R_{work}/R_{free}$ | 0.203/0.226 (0.271/0.307) |
| No. of atoms | |
| Protein | 4683 |
| Ion | 6 |
| Water | 226 |
| *B*-factors | |
| Protein | 45.8 |
| Ion | 31.5 |
| Water | 45.6 |
| rms deviations | |
| Bond lengths (Å) | 0.007 |
| Bond angles (º) | 1.15 |
| Ramachandran Plot (%)[1] | 97.9 / 2.1 / 0 |

$R_{free}$ was calculated with 5% of the data.

Numbers in parentheses represent values in the highest-resolution shell.

[1]Residues in favored, allowed, and disallowed regions of the Ramachandran plot.

**Table S2. $V_{1/2}$, $\Delta V_{1/2}$, and $z$ for 0, 100 μM and 10 mM $Ca^{2+}$ for WT and mutant channels**

| [Ca] | Channel | $V_{1/2}$ (mV) | $z$ |
|---|---|---|---|
| 0 μM | WT BK | 176 ± 0.9 | 1.06 ± 0.03 |
| | L385P[a] | 248 ± 1.6 | 0.98 ± 0.02 |
| | L387P[a] | 238 ± 4.0 | 0.80 ± 0.04 |
| | E388P[a] | 242 ± 3.9 | 0.77 ± 0.03 |
| | A389P[a] | 242 ± 3.4 | 0.90 ± 0.04 |
| | L390P[a] | 242 ± 2.5 | 0.93 ± 0.03 |
| | F391P[a] | 220 ± 2.3 | 0.78 ± 0.03 |
| | H394P[a] | 285 ± 2.8 | 0.98 ± 0.04 |
| | L390P/H394P[a] | 208 ± 2.5 | 0.90 ± 0.04 |
| 100 μM | WT BK | -32 ± 2.1 | 0.96 ± 0.07 |
| | L385P | 78 ± 0.9 | 0.97 ± 0.03 |
| | L387P | 135 ± 1.8 | 0.85 ± 0.03 |
| | E388P | 134 ± 1.1 | 0.84 ± 0.03 |
| | A389P | 121 ± 0.7 | 0.96 ± 0.02 |
| | L390P | 140 ± 0.6 | 0.95 ± 0.02 |
| | F391P | 140 ± 0.7 | 0.83 ± 0.02 |
| | H394P | 114 ± 0.9 | 1.03 ± 0.03 |
| | L390P/H394P | 157 ± 2.7 | 0.75 ± 0.05 |
| 10 mM | WT BK | -89 ± 2.3 | 0.98 ± 0.06 |
| | L387P | 93 ± 2.1 | 1.30 ± 0.12 |
| | E388P | 92 ± 1.8 | 1.15 ± 0.03 |
| | A389P | 74 ± 1.3 | 0.96 ± 0.08 |
| | F391P | 97 ± 1.9 | 1.17 ± 0.08 |

| [Ca] | Channel | $\Delta V_{1/2}$ (mV) | $\Delta V_{1/2}$ % of WT |
|---|---|---|---|
| 0 → 100 μM | WT BK | -208 ± 2.3 | 100 |
| | L385P[b] | -170 ± 1.8 | 82 |
| | L387P[b] | -102 ± 4.4 | 50 |
| | E388P[b] | -108 ± 4.1 | 52 |
| | A389P[b] | -120 ± 3.5 | 58 |
| | L390P[b] | -101 ± 2.6 | 49 |
| | F391P[b] | -80 ± 2.4 | 38 |
| | H394P[b] | -170 ± 2.9 | 82 |
| | L390P/H394P[b] | -51 ± 3.7 | 25 |
| 100 μM → 10 mM | WT BK | -58 ± 3.1 | 100 |
| | L387P[c] | -43 ± 2.8 | 74 |
| | E388P[c] | -43 ± 2.1 | 74 |
| | A389P[c] | -47 ± 1.5 | 82 |
| | F391P[c] | -43 ± 2.0 | 75 |

$V_{1/2}$ is the voltage where G/Gmax in the Boltzmann fits reaches 1/2 of the maximal activation and $z$ is effective gating charge (see Eqns. 1 and 2 in Materials and Methods).
$\Delta V_{1/2}$ = $V_{1/2}$ in 100 μM $Ca^{2+}$ - $V_{1/2}$ in 0 $Ca^{2+}$, and $V_{1/2}$ in 10 mM $Ca^{2+}$ - $V_{1/2}$ in 100 μM $Ca^{2+}$.
$\Delta V_{1/2}$ % of WT = ($\Delta V_{1/2}$ mutant / $\Delta V_{1/2}$ WT) x 100%.
[a]$V_{1/2}$ of mutant is > WT, $p < 0.00001$.
[b]Left shift of mutant is < WT, $p < 0.00001$.
[c]Left shift of mutant is < WT, $p < 0.03$.

**Table S3. $V_{1/2}$, $\Delta V_{1/2}$, and z for 0 and 100 μM $Ca^{2+}$ for WT and mutant channels**

| [Ca] | Channel | $V_{1/2}$ (mV) | *z* |
|---|---|---|---|
| 0 μM | WT BK | 176 ± 0.9 | 1.0 ± 0.03 |
| | 2D2A[a] | 188 ± 2.3 | 0.9 ± 0.05 |
| | 5D5N[b] | 225 ± 1.2 | 1.0 ± 0.02 |
| | 2D2A/5D5N[b] | 216 ± 2.0 | 0.9 ± 0.04 |
| | L390P/2D2A[b] | 248 ± 6.3 | 0.8 ± 0.05 |
| | L390P/5D5N[b] | 241 ± 3.6 | 1.0 ± 0.06 |
| | L390P/2D2A/5D5N[b] | 263 ± 6.3 | 0.7 ± 0.01 |
| 100 μM | WT | -32 ± 2.1 | 0.9 ± 0.07 |
| | 2D2A | 68 ± 1.8 | 0.8 ± 0.05 |
| | 5D5N | 58 ± 1.9 | 0.9 ± 0.06 |
| | 2D2A/5D5N | 200 ± 2.3 | 0.8 ± 0.04 |
| | L390P/2D2A | 183 ± 1.6 | 0.7 ± 0.02 |
| | L390P/5D5N | 186 ± 0.7 | 0.8 ± 0.02 |
| | L390P/2D2A/5D5N | 245 ± 2.3 | 0.7 ± 0.02 |
| [Ca] | Channel | $\Delta V_{1/2}$ (mV) | $\Delta V_{1/2}$ % of WT |
| 0 → 100 μM | WT | -208 ± 2.3 | 100 |
| | 2D2A[c,d,e] | -119 ± 2.9 | 58 |
| | 5D5N[c,d,e] | -166 ± 2.5 | 80 |
| | 2D2A/5D5N[c] | -15 ± 3.1 | 7 |
| | L390P/2D2A[c,e] | -64 ± 6.5 | 31 |
| | L390P/5D5N[c,e] | -54 ± 3.7 | 26 |
| | L390P/2D2A/5D5N[c] | -18 ± 6.7 | 9 |

$V_{1/2}$ is the voltage where G/Gmax in the Boltzmann fits reaches 1/2 of the maximal activation, and *z* is effective gating charge (see Eqns. 1 and 2 in Materials and Methods). 2D2A indicates BK channels with the RCK1 $Ca^{2+}$ sites disabled using mutations D362A/D367A. 5D5N indicates a BK channel with the $Ca^{2+}$ bowl sites disabled using mutation D898-902N.

$\Delta V_{1/2}$ = $V_{1/2}$ in 100 μM $Ca^{2+}$ - $V_{1/2}$ in 0 $Ca^{2+}$.

$\Delta V_{1/2}$ % of WT = ($\Delta V_{1/2}$ mutant / $\Delta V_{1/2}$ WT) x 100%.

[a]$V_{1/2}$ of mutant > WT, $p < 0.003$.

[b]$V_{1/2}$ of mutant > WT, $p < 0.00001$.

[c]Left shift of mutant < WT, $p < 0.00001$.

[d]Left shift for 2D2A < 5D5N, $p < 0.0001$.

[e]L390P reduced the left shift of 2D2A and 5D5N, $p < 0.0003$.

**Table S4. $V_{1/2}$, $z$, and $\Delta V_{1/2}$ for 0 and 10 mM $Mg^{2+}$ for WT and mutant channels**

| [Mg] | Channel | $V_{1/2}$ (mV) | $z$ |
|---|---|---|---|
| 0 mM | WT BK | 176 ± 0.9 | 1.06 ± 0.03 |
| | L385P | 248 ± 1.6 | 0.98 ± 0.02 |
| | L387P | 238 ± 4.0 | 0.80 ± 0.04 |
| | E388P | 242 ± 3.9 | 0.77 ± 0.03 |
| | A389P | 242 ± 3.4 | 0.90 ± 0.04 |
| | L390P | 242 ± 2.5 | 0.93 ± 0.03 |
| | F391P | 220 ± 2.3 | 0.78 ± 0.03 |
| | H394P | 285 ± 2.8 | 0.98 ± 0.04 |
| 10 mM Mg | WT BK | 116 ± 0.8 | 1.18 ± 0.04 |
| | L385P | 193 ± 0.6 | 1.04 ± 0.02 |
| | L387P | 202 ± 1.0 | 0.98 ± 0.03 |
| | E388P | 202 ± 0.7 | 0.94 ± 0.02 |
| | A389P | 194 ± 0.6 | 0.84 ± 0.01 |
| | L390P | 199 ± 0.4 | 1.05 ± 0.01 |
| | F391P | 186 ± 0.4 | 0.89 ± 0.01 |
| | H394P | 251 ± 1.0 | 1.07 ± 0.03 |
| **[Mg]** | **Channel** | **$\Delta V_{1/2}$ (mV)** | **$\Delta V_{1/2}$ % of WT** |
| 0 → 10 mM | WT BK | -60 ± 1.2 | 100 |
| | L385P[a] | -55 ± 1.7 | 92 |
| | L387P[b] | -36 ± 4.1 | 60 |
| | E388P[b] | -40 ± 4.0 | 67 |
| | A389P[c] | -48 ± 3.5 | 80 |
| | L390P[b] | -43 ± 2.5 | 72 |
| | F391P[d] | -34 ± 2.3 | 57 |
| | H394P[d] | -34 ± 3.0 | 57 |

$V_{1/2}$ is the voltage where G/Gmax in the Boltzmann fits reaches 1/2 of the maximal activation, and $z$ is effective gating charge (see Eqns. 1 and 2 in Materials and Methods). $\Delta V_{1/2} = V_{1/2}$ in 10 mM $Mg^{2+}$ - $V_{1/2}$ in 0 $Mg^{2+}$.

$\Delta V_{1/2}$ % of WT = ($\Delta V_{1/2}$ mutant / $\Delta V_{1/2}$ WT) x 100%.

[a]Left shift not significantly different from WT, $p > 0.05$.

[b]Left shift < WT, $p < 0.01$.

[c]Left shift < WT, $p < 0.05$.

[d]Left shift < WT, $p < 0.002$.

**References for Supporting Information**


1. P. Yuan, M. D. Leonetti, A. R. Pico, Y. Hsiung, R. MacKinnon, Structure of the human BK channel $Ca^{2+}$-activation apparatus at 3.0 A resolution. *Science* **329**, 182-186 (2010).
2. Z. Otwinowski, W. Minor, Processing of X-ray diffraction data collected in oscillation mode. *Methods Enzymol* **276**, 307-326 (1997).
3. A. J. Mccoy *et al.*, Phaser crystallographic software. *J Appl Crystallogr* **40**, 658-674 (2007).
4. P. Emsley, K. Cowtan, Coot: model-building tools for molecular graphics. *Acta Crystallogr D Biol Crystallogr* **60**, 2126-2132 (2004).
5. G. N. Murshudov, A. A. Vagin, E. J. Dodson, Refinement of macromolecular structures by the maximum-likelihood method. *Acta Crystallogr D Biol Crystallogr* **53**, 240-255 (1997) .
6. R. K. Hite, X. Tao, R. MacKinnon, Structural basis for gating the high-conductance $Ca^{2+}$-activated $K^+$ channel. *Nature* **541**, 52-57 (2017).
7. X. Tao, R. K. Hite, R. MacKinnon, Cryo-EM structure of the open high-conductance $Ca^{2+}$-activated $K^+$ channel. *Nature* **541**, 46-51 (2017).
8. N. Collaborative Computational Project, The CCP4 suite: programs for protein crystallography. *Acta Crystallogr D Biol Crystallogr* **50**, 760-763 (1994).
9. A. Butler, S. Tsunoda, D. P. McCobb, A. Wei, L. Salkoff, mSlo, a complex mouse gene encoding "maxi" calcium-activated potassium channels. *Science* **261**, 221-224 (1993).
10. G. Zhang *et al.*, Deletion of cytosolic gating ring decreases gate and voltage sensor coupling in BK channels. *J. Gen. Physiol* **149**, 373-387 (2017).
11. B. S. Rothberg, K. L. Magleby, Voltage and $Ca^{2+}$ activation of single large-conductance $Ca^{2+}$-activated $K^+$ channels described by a two-tiered allosteric gating mechanism. *J. Gen. Physiol* **116**, 75-99 (2000).
12. J. Cui, D. H. Cox, R. W. Aldrich, Intrinsic voltage dependence and $Ca^{2+}$ regulation of mslo large conductance Ca-activated $K^+$ channels. *J. Gen. Physiol* **109**, 647-673 (1997).
13. F. T. Horrigan, J. Cui, R. W. Aldrich, Allosteric voltage gating of potassium channels I. Mslo ionic currents in the absence of $Ca^{2+}$. *J. Gen. Physiol* **114**, 277-304 (1999).
